# Integrating patterns of thermal tolerance and phenotypic plasticity with population genetics to improve understanding of vulnerability to warming in a widespread copepod

**DOI:** 10.1101/619775

**Authors:** Matthew C. Sasaki, Hans G. Dam

## Abstract

Differences in population vulnerability to warming are defined by spatial patterns in thermal adaptation. These patterns may be driven by natural selection over spatial environmental gradients, but can also be shaped by gene flow, especially in marine taxa with high dispersal potential. Understanding and predicting organismal responses to warming requires disentangling the opposing effects of selection and gene flow. We begin by documenting genetic divergence of thermal tolerance and developmental phenotypic plasticity. Ten populations of the widespread copepod *Acartia tonsa* were collected from sites across a large thermal gradient, ranging from the Florida Keys to Northern New Brunswick, Canada (spanning over 20 degrees latitude). Thermal performance curves from common garden experiments revealed local adaptation at the sampling range extremes, with thermal tolerance increasing at low latitudes and decreasing at high latitudes. The opposite pattern was observed in phenotypic plasticity, which was strongest at high latitudes. Over a large portion of the sampled range, however, we observed a remarkable lack of differentiation of thermal performance curves. To examine whether this lack of divergence is the result of selection for a generalist performance curve or constraint by gene flow, we analyzed cytochrome oxidase I mtDNA sequences, which revealed abundant genetic diversity and widely-distributed haplotypes. Strong divergence in thermal performance within genetic clades, however, suggests that the pace of thermal adaptation can be relatively rapid. The combined insight from the laboratory physiological experiments and genetic data indicate that gene flow constrains differentiation of thermal performance curves. This balance between gene flow and selection has implications for patterns of vulnerability to warming. Taking both genetic differentiation and phenotypic plasticity into account, our results suggest that local adaptation does not increase vulnerability to warming, and that low latitude populations in general may be more vulnerable to predicted temperature change over the next century.

## Introduction

Temperature affects processes at every level of biological organization (Angilletta, 2009; Hochachka & Somero, 2002). The rapid warming of the world’s oceans (Cheng *et al*., 2019a; Cheng *et al*., 2019b; Roemmich *et al*., 2015; Saba *et al*., 2016; Wijffels *et al*., 2016) presents a significant threat to contemporary marine biodiversity (Bryndum-Buchholz *et al*., 2019; Parmesan, 2006). In addition to the increase in mean ocean temperature, significant increases in the magnitude and frequency of acute events like heat waves have been predicted (Lorenzo & Mantua, 2016; Meehl, 2004; Perkins *et al*., 2012). These acute events have significant consequences for organisms, and therefore cannot be ignored in predictions of biotic response to climate change (Campbell-Staton *et al*., 2017; Leicht *et al*., 2017; Stoks *et al*., 2017; Sydeman *et al*., 2013; Ummenhofer & Meehl, 2017; Wernberg *et al*., 2016). Vulnerability to these climatic changes is established by pre-existing spatial patterns in thermal adaptation. Characterizing patterns of thermal adaptation and determining their underlying causes is, therefore, directly related to our ability to predict vulnerability and responses of the biota to climate change (Moran *et al*., 2016; Sorte *et al*., 2011).

Macrophysiology, the study of variation in physiological traits across space and time (Chown *et al*., 2004), often yields evidence for adaptation across environmental gradients. Latitudinal thermal gradients, for example, are well-known drivers of local adaptation of thermal tolerance (Addo-Bediako *et al*., 2000; Castañeda *et al*., 2015; Gaitán-Espitia *et al*., 2017; Pereira *et al*., 2017; Yampolsky *et al*., 2013). These patterns in adaptation across large spatial scales are often attributed to selection acting on a set of populations, as gene flow is assumed to be relatively low over large distances. This important assumption may not always hold for marine taxa which often have high dispersal potentials (Bowen *et al*., 2016; Carlton *et al*., 2017; Cowen *et al*., 2006; Cowen & Sponaugle, 2009; Gélin *et al*., 2017; Kinlan & Gaines, 2003; Sexton & Norris, 2008). However, dispersal dynamics in the ocean can be complex (McManus & Woodson, 2012), preventing easy generalization or prediction of connectivity. Instead, genetic markers are often required to estimate levels of connectivity between populations (Palumbi, 2003).

Adaptive genetic differentiation and phenotypic plasticity are two of the main mechanisms used by organisms to cope with variation in the thermal environment. (Angilletta, 2009; Dam, 2013; Magozzi & Calosi, 2014; Somero, 2010; Sparks *et al*., 2017). Adaptive genetic differentiation is well-known to produce significant variation in phenotypes (Hochachka & Somero, 2002). Phenotypic plasticity, the capacity of a single genotype to produce multiple phenotypes in response to different environmental conditions, can also have large effects (Ayrinhac *et al*., 2004; Chown *et al*., 2004, West-Eberhard 2003). Several types of phenotypic plasticity, including acclimation (Stillman, 2003), hardening (Sørensen *et al*., 2001), and developmental phenotypic plasticity (Pereira *et al*., 2017), have all been shown to have strong effects on organismal thermal tolerance. Both mechanisms are likely to play important roles in determining organismal responses to climate change (Hoffmann & Sgro, 2011; Reusch, 2013). Importantly, plasticity acts within generations, and might therefore provide a mechanism for rapid response to environmental variability (Chevin *et al*., 2010; Chown *et al*., 2007; Merilä & Hendry, 2014; Seebacher & Grigaltchik, 2014). Plasticity may also prevent the loss of cryptic genetic diversity by shielding genotypes from selection (Friedrich & Meyer, 2016; Pfennig *et al*., 2010; Schlichting, 2004). This is in stark contrast to selection on standing genetic diversity, which may result in strong demographic bottlenecks and the loss of genetic diversity (Corbett-Detig *et al*., 2015; Hoffmann & Sgrò, 2011; Kellermann *et al*., 2009). Studies that examine both mechanisms across large spatial scales are needed.

Characterizing the spatial patterning of phenotypic plasticity and adaptive genetic differentiation is important for predictions of population vulnerability to warming. The Climate Variability Hypothesis (CVH) (Janzen, 1967; Stevens, 1989) posits that thermal tolerance should correspond to the mean temperature experienced by a population whereas phenotypic plasticity should evolve in response to variability in the thermal environment. This hypothesis has accumulated support over time, especially in terrestrial and freshwater systems (Deutsch *et al*., 2008; Sunday *et al*., 2010), but still lacks robust experimental validation in the marine realm. It has also been proposed that patterns in the evolution of phenotypic plasticity may be the result of a trade-off between thermal tolerance and the strength of phenotypic plasticity (Stillman, 2003), where higher thermal tolerance evolves at the expense of the capability to modify the phenotype via plasticity.

Patterns in adaptation can, however, also be strongly influenced by gene flow (Lenormand, 2002). The “Gene Flow vs. Selection” issue has been at the heart of evolutionary ecology for decades (Blanquert *et al*. 2013; Slatkin, 1985; 1987). Successful gene flow between populations can strongly impede local adaptation (Garant *et al*., 2007; Hendry & Taylor, 2004; Lenormand, 2002; Moore *et al*., 2007; Nosil & Crespi, 2004), and phenotypic divergence is often correlated with the degree of isolation (Mayr, 1963). However, low levels of gene flow might also promote local adaptation by increasing the genetic diversity contained within a population (Garant *et al*., 2007; Tallmon *et al*., 2004). The potential for interaction between selection and gene flow makes integrated approaches to studying evolutionary physiology critical for robust characterization of spatial patterns in adaptation. Taking both selection and gene flow into account may be needed to explain observed patterns of adaptation (Dionne *et al*., 2008; Moore & Hendry 2005).

Spatial patterns in both adaptation and predicted warming interact to produce what is likely to be spatially heterogeneous vulnerability to warming. Understanding which populations are more vulnerable to warming, and why, is critical for effective management and conservation of diversity. Previous work has suggested that warm-adapted, low latitude species or populations are more vulnerable to climate change as they already experiencing temperatures near their thermal maxima (Comte & Olden, 2017; Tewksbury *et al*., 2008; Vinagre *et al*., 2016). However, this is not a universal observation, and populations from mid- to high-latitudes have also been predicted to be more vulnerable (Bennett *et al*., 2015; Calosi *et al*., 2008; Fusi *et al*., 2015). Additionally, these predictions are often based on measurements of thermal tolerance. This is insufficient, however, as phenotypic plasticity may also play a large role in determining vulnerability to climate change (Burggren, 2018; Chown *et al*., 2007; Magozzi & Calosi, 2015; Sparks *et al*., 2017). Examining spatial patterns in both thermal tolerance and the strength of phenotypic plasticity may provide more robust estimates of vulnerability.

Copepods are the most abundant metazoans in the ocean (Humes, 1994; Turner, 2004). They play an important role in transferring energy from primary producers to secondary consumers and are therefore tightly linked to global biogeochemical systems (Menden-Deuer & Kiørboe, 2016), marine trophic webs (Turner, 2004), and commercial fisheries (Castonguay *et al*., 2008; Dam and Baumann, 2017; Friedland *et al*., 2012). Studies of thermal adaptation in copepods have a long history (Bradley, 1978; Lonsdale & Levinton, 1985), but local adaptation has largely been ignored, especially in pelagic species (but see Smolina *et al*., 2016). The calanoid copepod *Acartia tonsa* often dominates coastal and estuarine environments. With a large latitudinal distribution (Turner, 1981) and a life-history amenable to laboratory culturing, this is an ideal model species for studying spatial patterns in adaptation. Previous studies of *Acartia* species have observed local adaptation to several different factors, including exposure to toxic dinoflagellates (Colin & Dam, 2002; 2004), salinity (Plough *et al*., 2018), and pH (Aguilera *et al*., 2016). Limited evidence also exists that local adaptation to temperature occurs across large geographic scales (González, 1974; Sasaki *et al*., 2019). This past work suggests that low latitude populations may be more vulnerable to warming (Sasaki *et al*., 2019), but conclusions are limited by the lack of spatial coverage and the number of populations used. Additionally, no population genetic information was included. Population genetic studies have shown genetic structuring in this taxon to be complex, with abundant cryptic diversity (Caudill & Bucklin, 2004; Chen & Hare, 2011). The effects of this genetic diversity on patterns of adaptation has largely been ignored (but see Plough *et al*. 2018). Here we examine both large- and fine-scale spatial patterns in the effects of genetic differentiation and phenotypic plasticity on the thermal performance curves (TPCs) of *Acartia tonsa*. We observe clear latitudinal variation in both thermal tolerance and the magnitude of the plastic response, but also find a remarkable lack of divergence over a large portion of the sampled range. Paired with insights from the analysis of cytochrome oxidase I (COI) mtDNA sequence data, we suggest that this lack of divergence is due to constraint by gene flow rather than selection for a generalist performance curve. These spatial patterns in adaptation will likely affect vulnerability to warming, with southern populations being the most vulnerable.

## Methods

### Sampling and Culture Maintenance

Copepods were collected from ten sites spanning 21 degrees of latitude, ranging from the Florida Keys to the Northumberland Strait in Canada (Fig. 1; Table 1). Sites selected cover a range of thermal environments, with mean monthly temperatures ranging from 9.05 – 25.1°C, and average monthly temperature ranges varying between 1.38 – 12.88°C. At each site, samples were collected in surface plankton tows using a 250-μm mesh plankton net with a solid cod end. Water column depth was less than 10 m at all sites. Salinity and surface water temperature were recorded at the time of collection. Within 3 hours of collection, mature *Acartia tonsa* individuals were visually identified using a dissection microscope and sorted into 0.6-μm filtered sea water (FSW), with salinity and temperature adjusted to match collection conditions. Each culture began with more than 1000 mature females and abundant males to ensure fertilization. Additional individuals were preserved in 95% molecular grade ethanol for later genetic analysis. All samples and cultures were then transported by car back to the University of Connecticut, Avery Point campus. Copepods were transported in temperature-controlled containers with particular care to maintain temperature and salinity near collection conditions. Aquarium bubblers were used to keep containers well oxygenated. Copepods were fed with a mixture of a green flagellate (*Tetraselmis* sp.) and a small diatom (*Thalassiosira weissflogii*) during transport. In the laboratory, live cultures were gradually brought to 18°C and 30 practical salinity units (psu) and then maintained for several generations under constant conditions to minimize the effects of previous environmental acclimation. Aquarium bubblers were used to ensure cultures were well oxygenated. The water of each culture was changed weekly. During this period, cultures were fed *ad libitum* a mixture of a green flagellate (*Tetraselmis* sp.), a small diatom (*Thalassiosira weissflogii*), and a cryptomonad (*Rhodomonas salina*). Phytoplankton were cultured semi-continuously in F/2 medium (without silica for *Tetraselmis* and *Rhodomonas*) with a 12 hr:12 hr light:dark cycle at 18°C. Genetic samples were kept at −80°C until extraction.

**Figure 1:**
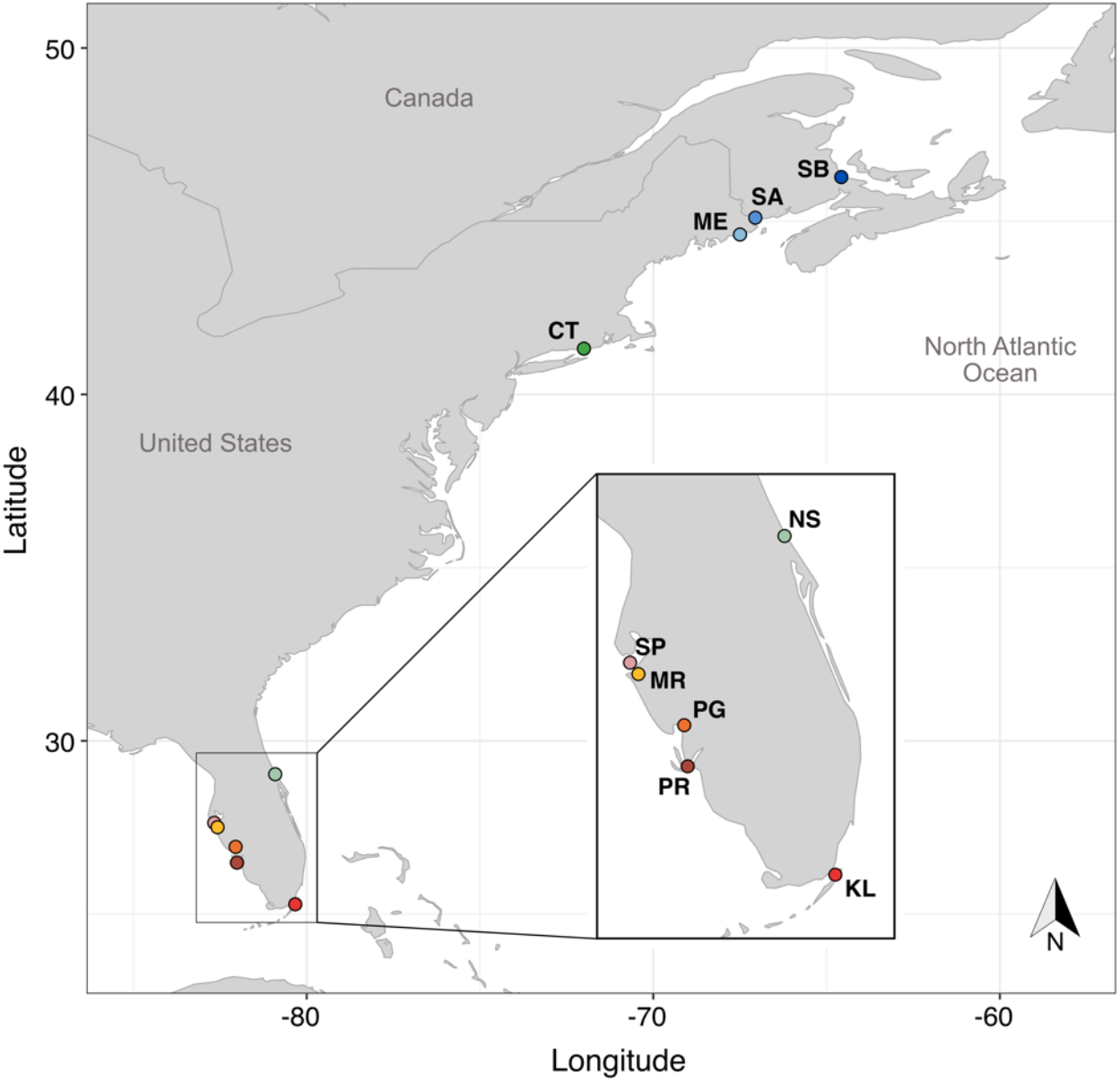
Map of sampling locations in the North Atlantic and Gulf of Mexico. The inset is a closer view of the southern sampling sites, to better show the spatial arrangement. Sampled site names are abbreviated: Shediac Bay (SB), St. Andrew (SA), Maine (ME), Connecticut (CT), New Smyrna (NS), St. Petersburg (SP), Manatee River (MR), Punta Gorda (PG), Punta Rasa (PR), and Key Largo (KL). Additional site details can be found in Table 1.

**Table 1:**
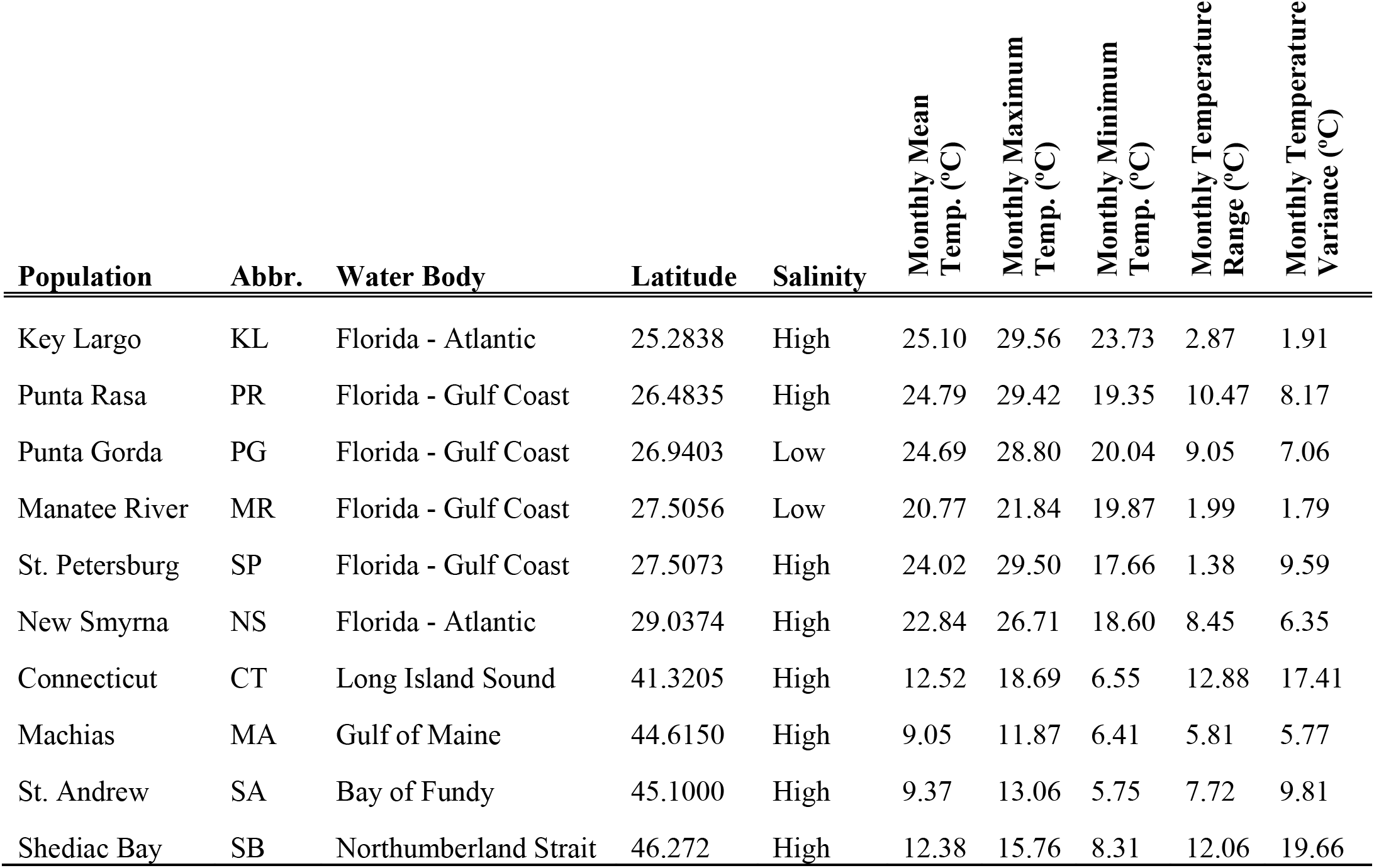
Collection site details for each of the ten populations recovered. Site names are provided, along with the two-letter abbreviation used. Water body refers to the general area where each site is located. The latitude of each site is provided, along with the salinity of collection. High salinity refers to sites with a salinity greater than 28 psu, while the low salinity sites both had a salinity of 5 psu. Climatological temperature data for each of the sites are also included. Raw temperature data was acquired from the MODIS-aqua sea surface temperature database and summarized to monthly intervals using CDO (Schulzweida *et al*., 2009).

### Heat Stresses and Performance Curve Estimation

After two generations, a split-brood common garden experiment was used to examine the effects of genetic differentiation and developmental phenotypic plasticity on the thermal performance curve (TPC) of *Acartia tonsa*. For each population, eggs were collected and randomly split between two developmental temperatures, 18°C and 22°C. All other conditions were kept the same between treatments. Upon reaching maturity, females were collected from both developmental conditions and exposed to a 24-hour acute heat stress. Healthy individuals were placed into 1.5 mL of FSW in a 2.0 mL microfuge tube, which was then partially capped to allow for gas exchange with the atmosphere. Only one female was placed into each tube. After all tubes were filled, they were left to rest for one hour at their respective developmental temperatures. Tubes were then placed into dry baths and exposed to a single temperature, ranging from 18°C to 38°C. The number of females per temperature varied, with fewer individuals exposed to the lowest and highest temperatures (where survivorship was expected to be least variable) and more individuals at intermediate temperatures. At least six females were used per temperature. Each female experienced only one temperature throughout the duration of the assay and was used for only one assay. After 24 hours, tubes were collected, and survivorship was determined visually using a dissection scope. Copepods were marked as alive if there was an active response to external stimuli or visible gut passage movement. Evaporation (and therefore fluctuations in salinity) was negligible during the heat stress.

All statistical analyses were performed using the software package R (R Core Team, 2016). Logistic regressions were used to estimate TPCs for the two developmental treatments for all ten populations. A three-way ANOVA was used to examine the effects of genetic differentiation and phenotypic plasticity on the TPCs (survival as a function of stress temperature, population, and developmental temperature). An effect of genetic differentiation would be indicated by a significant population term, while an effect of developmental phenotypic plasticity would be indicated by a significant developmental temperature term. Population differences in the strength of phenotypic plasticity would be indicated by a significant population x developmental temperature term (a heterogeneity of slopes test).

Each curve was also summarized by estimation of LD_50_ values (the temperature at which 50% survivorship would be observed). This is a common metric for thermal tolerance. We define the difference in LD50 values between the two developmental treatments (ΔLD_50_) as the strength of developmental phenotypic plasticity. Standard error for ΔLD_50_ values were calculated as ⎷(SE_18C_^2^ + SE_22C_^2^), where SE_18C_ and SE_22C_ are the standard error estimates for LD_50_ from the 18°C and 22°C developmental temperature groups, respectively. The effect of plasticity was also estimated by calculating the average increase at all survivorship levels from 10% - 90%, which we call ΔLD_average_. This takes into account any potential population-specific non-linearities in the effect of phenotypic plasticity across the TPC. We also estimated LD_10_ (temperature of 10% survival) and ΔLD_10_ values as estimates of thermal limits.

### Climatologies and Correlations

Daily satellite sea-surface temperature (SST) measurements for the years 2000 - 2017 were retrieved for each geographic location from the MODIS-Aqua database (https://oceancolor.gsfc.nasa.gov/data/aqua/). Because *A. tonsa* has generation times on the order of weeks to a month, this data was summarized as average monthly climatological parameters (mean, maximum, and minimum temperatures as well as temperature range and variance; Table 1) using the command line Climate Data Operators (CDO; Schulzweida *et al*., 2009).

The CVH postulates that thermal tolerance (LD_50_) should respond to some mean representation of the thermal environment, while the strength of the phenotypic plasticity (ΔLD_50_) should reflect the variation in the thermal environment. To test these predictions, we regressed our metrics of thermal adaptation against the environmental parameters of interest (LD_50_ ~ developmental temperature + monthly mean + monthly maximum + monthly minimum; ΔLD_50_ ~ monthly temperature variance + monthly temperature range). A linear regression was also used to examine the relationship between thermal tolerance and the strength of plasticity. Trade-offs between thermal tolerance and the strength of phenotypic plastic (Stillman, 2003) would be suggested by a significant negative relationship between LD_50_ and ΔLD_50_. Correlations between the environmental parameters and latitude, as well as with each other were also examined.

### Vulnerability to Warming

The statistical model used to estimate TPCs from the common garden experiment can also be used to predict what would happen if parameters like developmental temperature and stress temperature change. We use this to estimate vulnerability to warming across the ten populations examined. For each population, we estimated the TPC with the mean monthly temperature as the developmental temperature, using the *predict.glm* function in R (*stats* package). Survivorship at the mean monthly maximum temperature was then taken from this modelled TPC. The magnitude of predicted warming at each site over the next century was visually estimated from a high resolution model of warming in the North Atlantic (Saba *et al*. 2016). Future survivorship was estimated in a similar manner using these future temperature values (Supp. Table 1). Vulnerability was estimated as the difference between current and future survivorship values; a positive difference indicates an increase in survivorship (less vulnerable to warming) while a negative value indicates a decrease in survivorship (more vulnerable to warming).

### DNA Extraction, Amplification, and Sequencing

Mitochondrial cytochrome-oxidase I (COI) sequence data was generated for individuals from all ten populations (n = 17 - 34 per population). DNA was extracted using a Qiagen Blood and Tissue kit following the manufacturer’s instructions. Extracted DNA was eluted in 50 μl of elution buffer (25 μl twice) and stored at −20°C. COI sequences were amplified by polymerase chain reaction (PCR) using mtCOI primers LCO1490 (forward: GGTCAACAAATCATAAAGATATTGG) and HCO2198 (reverse: TAAACTTCAGGGTGACCAAAAAATCA) (Folmer *et al*., 1994). All PCR reactions were performed in 24 μl volumes with 13 μl ExTaq HS polymerase (Takara Bio Inc.), 1 μL each forward and reverse primers, 5 μl genomic DNA, and 5 μl ultrapure molecular grade water. The optimized PCR protocol began with an initial denaturation of 94°C for 3 minutes followed by 35 cycles of denaturation at 94°C for 45 seconds, annealing at 48°C for 45 seconds, and extension at 72°C for 45 seconds. The protocol ended with a final extension at 72°C for 7 minutes. Amplification success and product length were confirmed visually using a 1.2% agarose gel post-stained with GelRed (Biotium Inc.). Successful amplification products were then purified using an ExoSAP-IT PCR clean-up kit (ThermoFisher Scientific) following manufacturer’s instructions before being sent to Eurofins Genomics for forward and reverse strand sequencing.

### Sequence Analysis

Consensus sequences were generated for each individual using forward and reverse strands in UniPro (Okonechnikov *et al*., 2012). Sequences were aligned using Clustal-W (Thompson *et al*., 1994) and then visually checked. Species identity of each sequence was verified by BLAST search in NCBI’s GenBank database (Sayers *et al*., 2018). Population genetic summary statistics (nucleotide diversity (π), haplotype diversity (Hd), and average number of nucleotide differences between haplotypes) were calculated using DNaSP v6 (Librado & Rozas, 2009). A haplotype network using an infinite site model was computed using the R package *Pegas* (Paradis, 2010). MigrateN was used to estimate mutation-scaled population size and migration rate values (Beerli, 2006; Beerli & Felsenstein, 2001). The transition-transversion ratio was set to 20. A uniform migration rate prior was set with a minimum of 0 and a maximum of 2000, while the population size prior was set with a minimum of 0.001 and a maximum of 0.1. Analyses entailed a single long MCMC chain with 4 concurrent replicates and a static heating scheme. 10,000,000 steps were recorded with a burn-in of 500,000. All other settings used default values. The number of migrants per generation (N_M_) was then calculated as N_M(pop1->pop2)_ = Θ_pop1_ * M_pop1->pop2_. A linear regression between the number of migrants per generation and the pairwise population difference in LD_50_ was used to investigate constraint of thermal adaptation by gene flow.

## Results

### Differentiation of Thermal Performance Curves

A total of 6144 females were used to estimate the thermal performance curves (Fig. 2). The ANOVA results for the TPCs yield a significant effect of both population and developmental temperature (both p < 2.2×10^−16^; Table 2), indicating an influence of both genetic differentiation and developmental phenotypic plasticity, respectively. The significant interaction term between stress temperature and population (p < 2.2×10^−16^) indicates that genetic differentiation results in changes to the shape of the performance curve, while the significant interaction term between population and developmental temperature (p = 1.45×10^−7^) indicates that the strength of phenotypic plasticity differs between populations.

**Figure 2:**
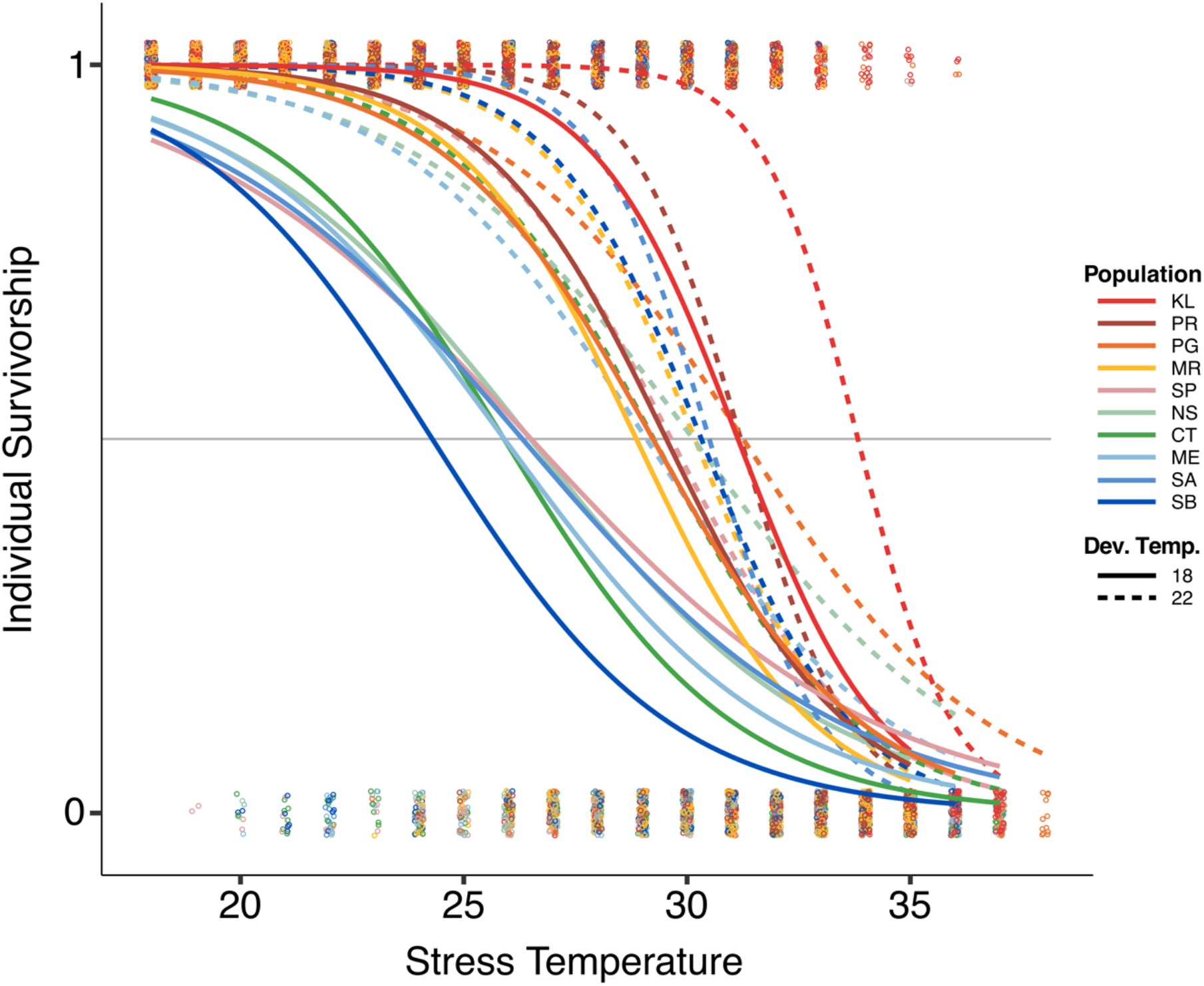
Thermal survivorship curves for each of the ten populations (different colors) and developmental temperatures (solid vs. dashed lines). Survivorship was measured using individuals from a split-brood, common garden experiment and a 24-hour acute heat shock. Survivorship was recorded as binary data (1 = survived, 0 = died), with curves estimated by logistic regression.

**Table 2:** Results of an ANOVA on the logistic regression of survivorship against stress temperature, population, and developmental temperature. All terms are statistically significant (*p* values ≪ 0.0002)

|  | **Df** | **Deviance** | **Resid. Df** | **Resid. Dev** | **Pr(>Chi)** |
| --- | --- | --- | --- | --- | --- |
| Stress Temp | 1 | 3024.26 | 6140 | 5158.1 | < 2.2e-16 |
| Population | 9 | 351.38 | 6131 | 4806.8 | < 2.2e-16 |
| Dev Temp | 1 | 266.91 | 6130 | 4539.8 | < 2.2e-16 |
| Stress Temp * Pop | 9 | 102.14 | 6121 | 4437.7 | < 2.2e-16 |
| Stress Temp * Dev Temp | 1 | 23.95 | 6120 | 4413.8 | 9.90E-07 |
| Pop * Dev Temp | 9 | 49.31 | 6111 | 4364.4 | 1.45E-07 |
| Stress Temp * Pop * Dev Temp | 9 | 32.45 | 6102 | 4332 | 0.0001665 |

These results are clearly evident in the TPCs (Fig. 2). Shediac Bay, the northernmost population, has a TPC shifted towards cooler temperatures while the southernmost population, Key Largo, has a TPC shifted towards warmer temperatures. Two groups of intermediate TPCs are also observed, one containing the populations from Manatee River, Punta Gorda, and Punta Rasa, with the other comprised of all remaining populations. This pattern can also be seen in the LD_50_ values of the 18°C developmental temperature group (Fig. 3). Differences are reduced in the 22°C developmental temperature group.

**Figure 3:**
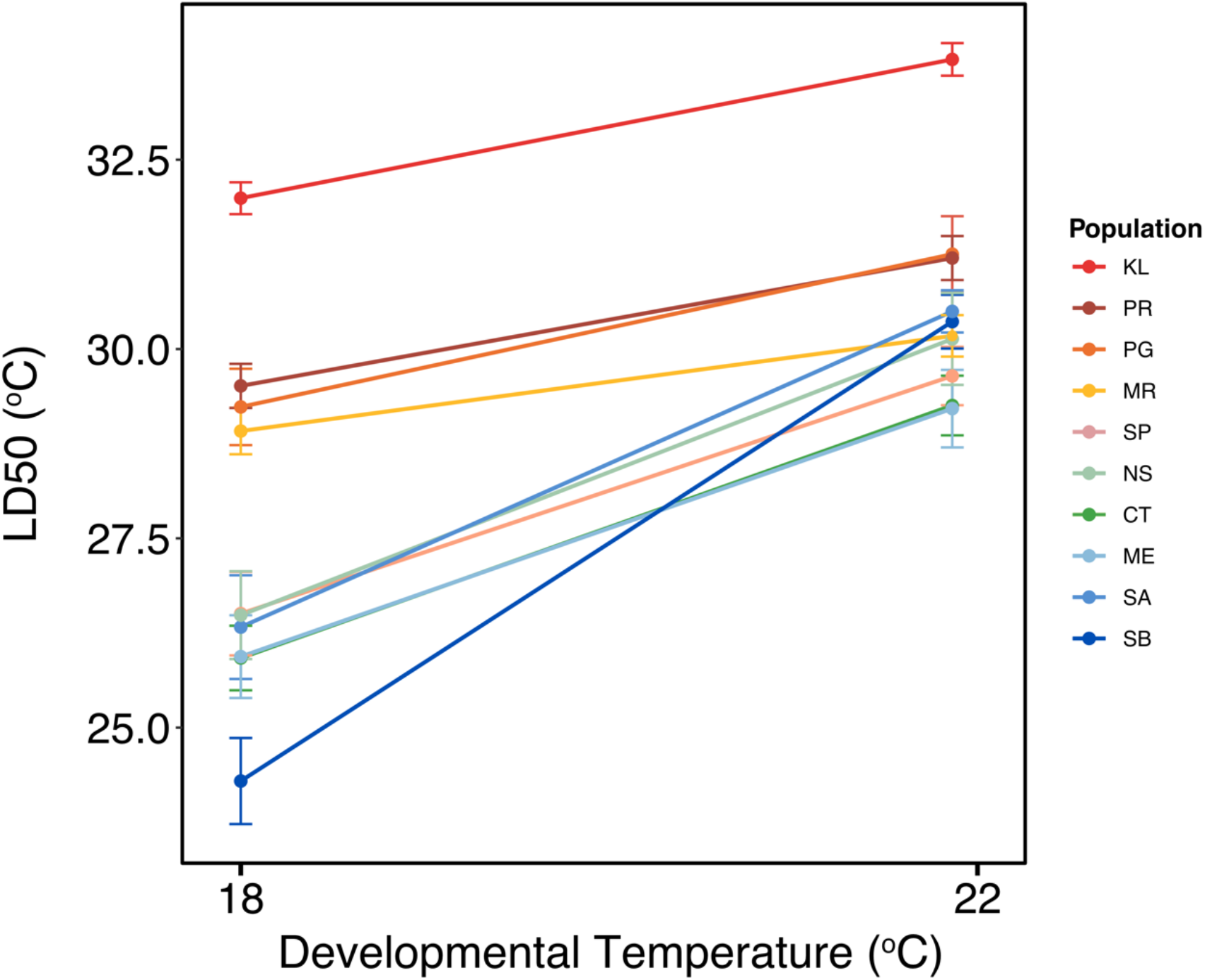
Reaction norms of thermal tolerance (LD_50_) for the ten populations, shown in different colors. LD_50_ was calculated as the temperature at which 50% survivorship was observed in the thermal performance curve. Error bars show standard errors. The slope of the individual norms represents the strength of developmental phenotypic plasticity (ΔLD_50_).

Thermal tolerance increased with warmer developmental temperature in all populations, suggesting a ubiquitous effect of developmental phenotypic plasticity. Shediac Bay, the northernmost population had the largest strength of phenotypic plasticity while Manatee River, a population from the Gulf Coast of Florida, had the smallest. The various crossed reaction norms also indicate variation in the strength of plasticity. The effects of developmental phenotypic plasticity were stronger for thermal tolerance than thermal limits (ΔLD_50_ > ΔLD_10_; Supp. Fig. 1). There are no large differences between ΔLD_50_ and ΔLD_average_ (Supp. Fig. 2), so we will focus only on ΔLD_50_ for the sake of uniformity with the LD_50_ metric of thermal tolerance.

### Lack of Differentiation in Thermal Performance Curves

Interestingly, we also observed a striking lack of differentiation in the TPCs of populations from a large portion of the sampling range, indicated by the boxes in Fig. 4. Thermal tolerance values for the St. Petersburg (SP), New Smyrna (NS), Connecticut (CT), Maine (ME), and Saint Andrew (SA) populations, spanning over 20 degrees latitude and originating from drastically different thermal environments, were remarkably similar. This lack of differentiation is also seen in the strength of developmental phenotypic plasticity.

**Figure 4:**
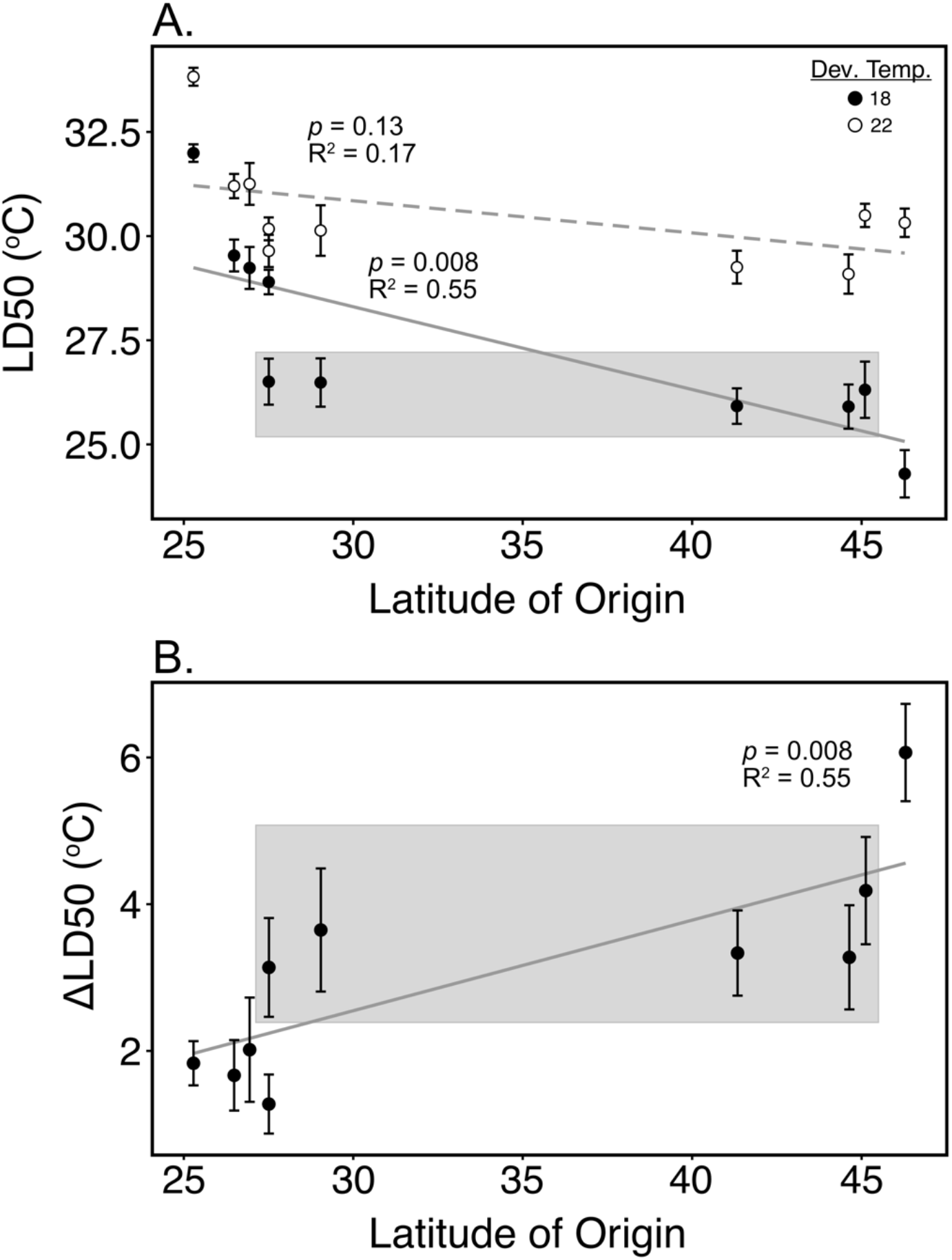
Latitudinal patterns in thermal adaptation. A) Thermal tolerance (LD_50_) is plotted against latitude of origin. Filled circles represent thermal tolerance values from the 18°C developmental temperature while unfilled circles are values from the 22°C developmental temperature. B) Developmental phenotypic plasticity (ΔLD_50_) is plotted against latitude of origin. In both panels, error bars represent standard error. Boxes highlight the latitudinal region with limited divergence of thermal performance curves.

### Environmental Correlations

Despite the lack of differentiation between some populations, LD_50_ and ΔLD_50_ were both significantly correlated with latitude (p = 0.0017 and 0.008 respectively; Table 3; Fig. 4). ANOVA results show LD_50_ to be significantly correlated with both mean monthly temperature (p = 3.7×10^−4^) and mean monthly minimum temperature (p = 0.0012; Table 4; Fig. 5). ΔLD_50_ is significantly correlated with mean monthly temperature variance (p = 0.015) but not mean monthly temperature range (p = 0.58; Table 4; Fig. 5). Both mean monthly temperature and mean monthly temperature variance were significantly correlated with latitude (p = 1.81×10^−6^ and 0.045 respectively; Supp. Fig. 3, 4) but not with each other (p = 0.14; Supp. Fig. 5). There is a significant correlation between LD_50_ and ΔLD_50_ (p = 0.0023; Fig. 6).

**Figure 5:**
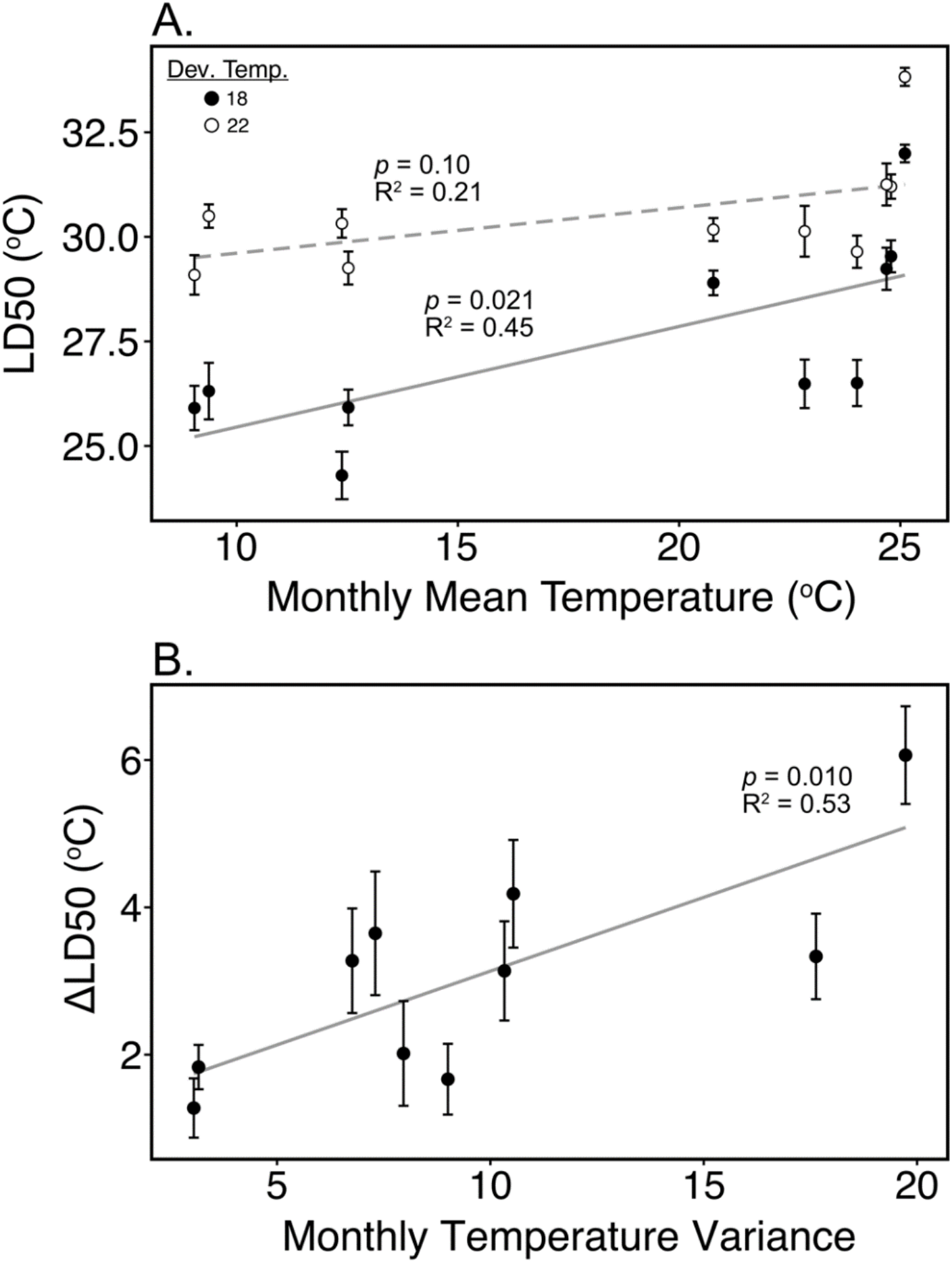
Correlation between thermal performance metrics and environmental parameters, as predicted by the Climate Variability Hypothesis. A) Thermal tolerance (LD_50_) is correlated with mean monthly temperature. Filled circles represent thermal tolerance values from the 18°C developmental temperature while empty circles are values from the 22°C developmental temperature. B) Phenotypic plasticity (ΔLD_50_) is correlated with mean monthly temperature variance. In both panels, error bars represent standard error.

**Figure 6:**
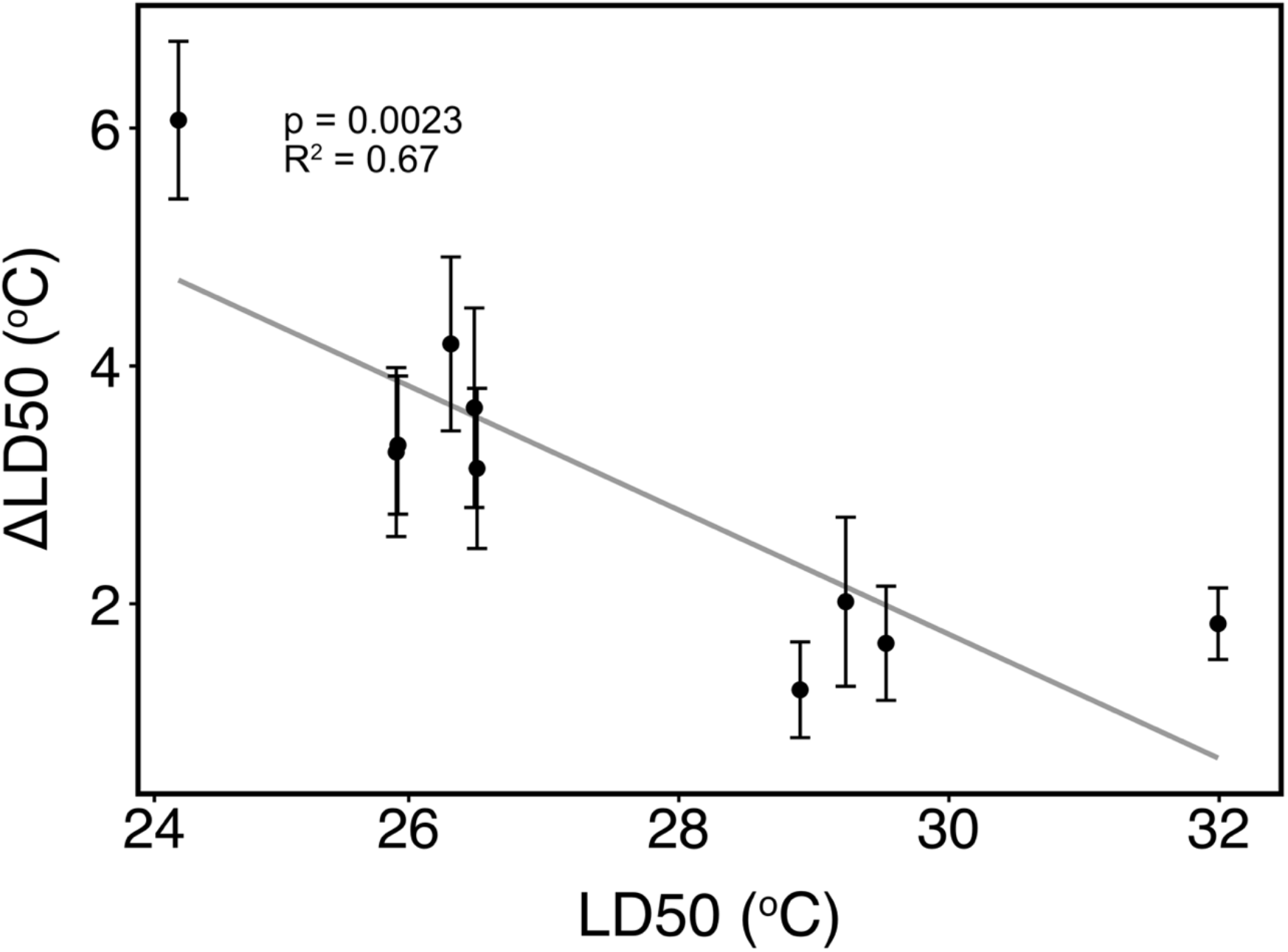
Linear correlation between thermal tolerance (LD_50_) and the strength of developmental phenotypic plasticity (ΔLD_50_) for the ten populations. Error bars represent standard error.

**Table 3:**
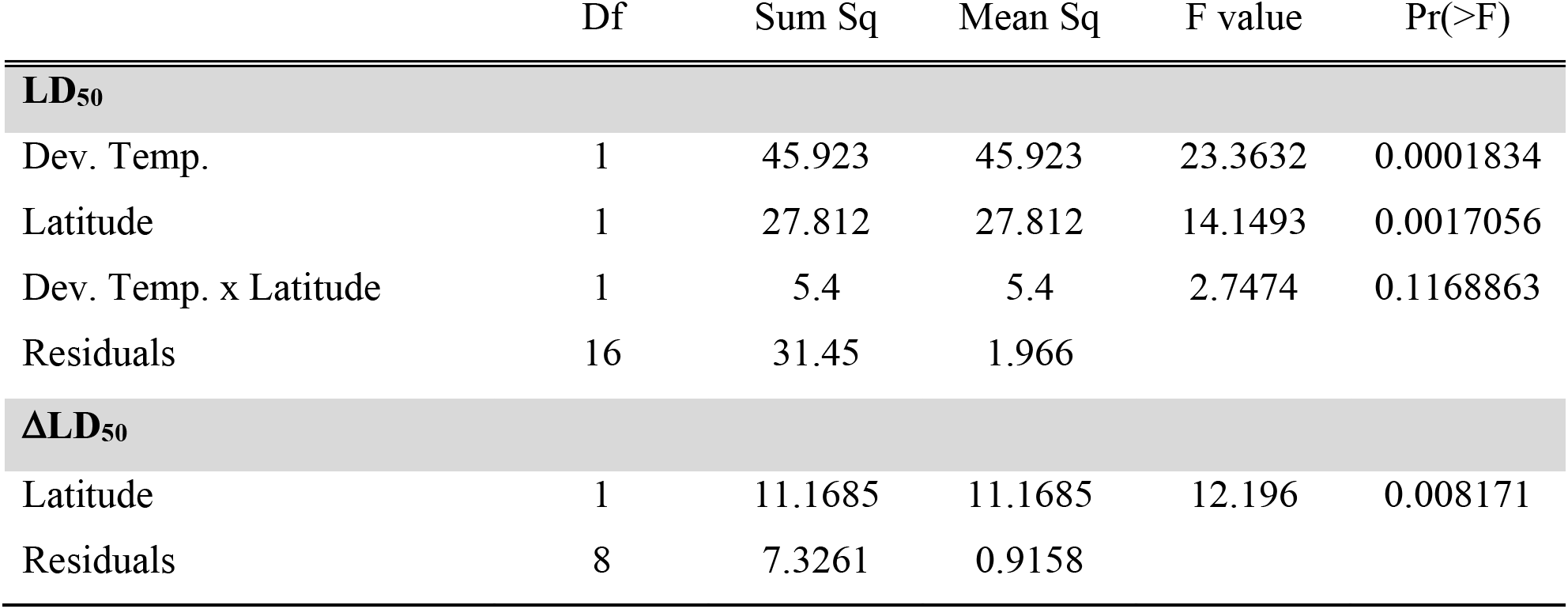
ANOVA results for the correlation between thermal tolerance and the strength of phenotypic plasticity with latitude.

**Table 4:** ANOVA results for the correlation between thermal tolerance and the strength of phenotypic plasticity with environmental parameters.

|  | Df | Sum Sq | Mean Sq | F value | Pr(>F) |
| --- | --- | --- | --- | --- | --- |
| **LD<sub>50</sub>** |  |  |  |  |  |
| Dev. Temp. | 1 | 45.923 | 45.923 | 37.1471 | 2.05E-05 |
| Monthly Mean | 1 | 25.722 | 25.722 | 20.8062 | 0.0003746 |
| Monthly Maximum | 1 | 0.693 | 0.693 | 0.5603 | 0.4657218 |
| Monthly Minimum | 1 | 19.704 | 19.704 | 15.9388 | 0.0011775 |
| Residuals | 15 | 18.544 | 1.236 |  |  |
| **ΔLD<sub>50</sub>** |  |  |  |  |  |
| Monthly Variance | 1 | 10.8028 | 10.8028 | 10.3074 | 0.01485 |
| Monthly Range | 1 | 0.3555 | 0.3555 | 0.3392 | 0.57859 |
| Residuals | 7 | 7.3364 | 1.0481 |  |  |

### Vulnerability to Warming

Change in survivorship at mean monthly maximum temperatures ranged from −16% to +9.8% (Supp. Table 1). There was a general trend in vulnerability across latitudes (Fig. 7). Locally adapted populations from low latitudes either saw almost no change (KL and MR) or a reduction in survivorship (PG and PR). The locally adapted population from high latitudes (SB) saw an increase in survivorship. Generally, populations without differentiation of TPCs saw a decrease in survivorship at low latitudes (NS and SP) but an increase in survivorship at high latitudes (CT and ME).

**Figure 7:**
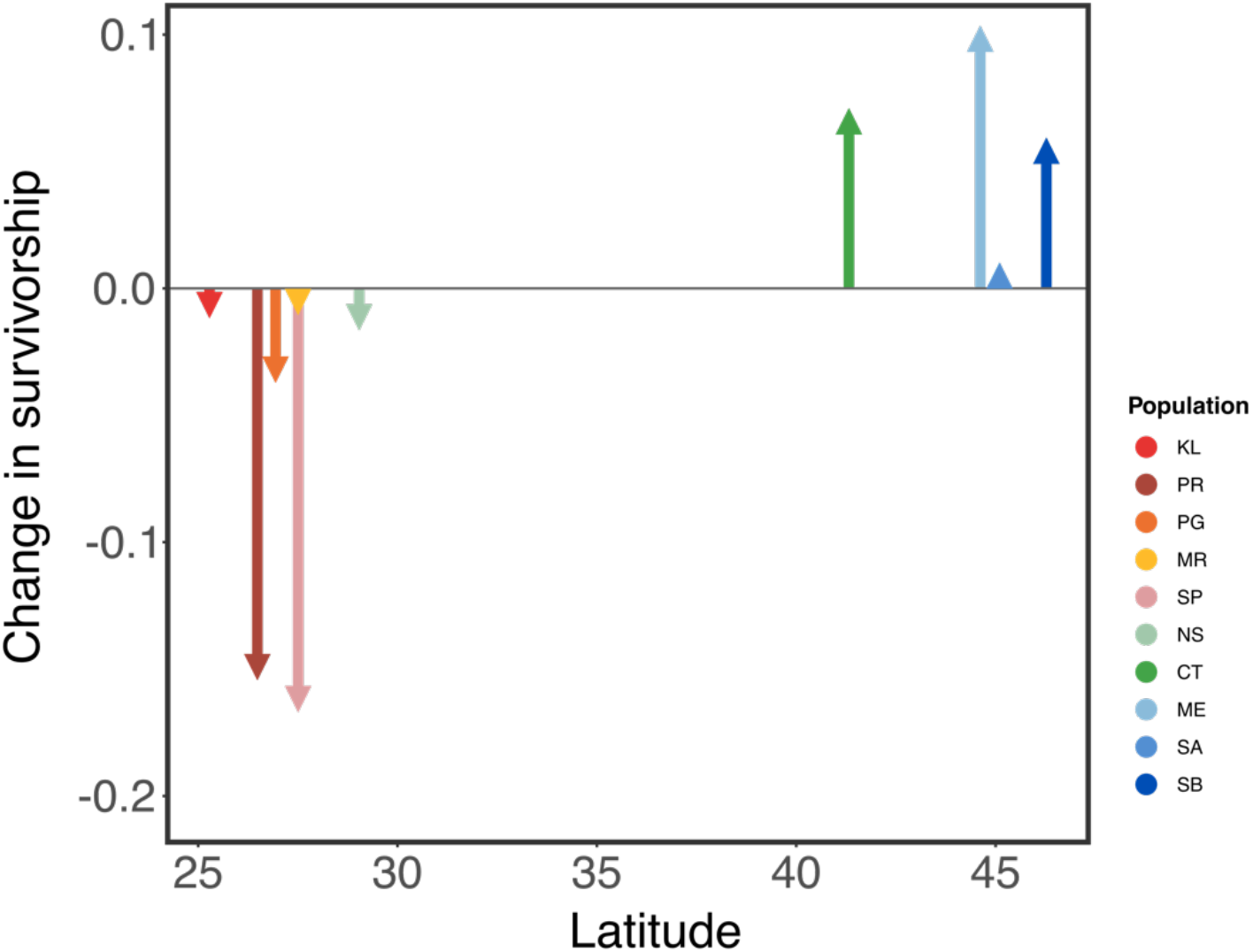
Vulnerability to predicted warming over the next century. Vulnerability is estimated as the change in survivorship at the mean monthly maximum temperature if the developmental temperature is assumed to be the mean monthly temperature. Positive changes represent an increase in survivorship while negative changes represent a decrease in survivorship. Estimates of the magnitude of warming and the temperature values used for estimating change in survivorship for each population can be found in Supp. Table 1.

### Genetic Diversity and Gene Flow

A total of 228 COI sequences (aligned length: 562 base pairs) were used for the genetic analyses. COI sequences revealed high levels of genetic diversity (Table 5). Large variation between populations was observed in haplotype diversity (Hd; 0.189 - 0.941), nucleotide diversity (π; 0.0029 - 0.079), and the average number of nucleotide differences between the haplotypes (1.47 - 40.46).

**Table 5:**
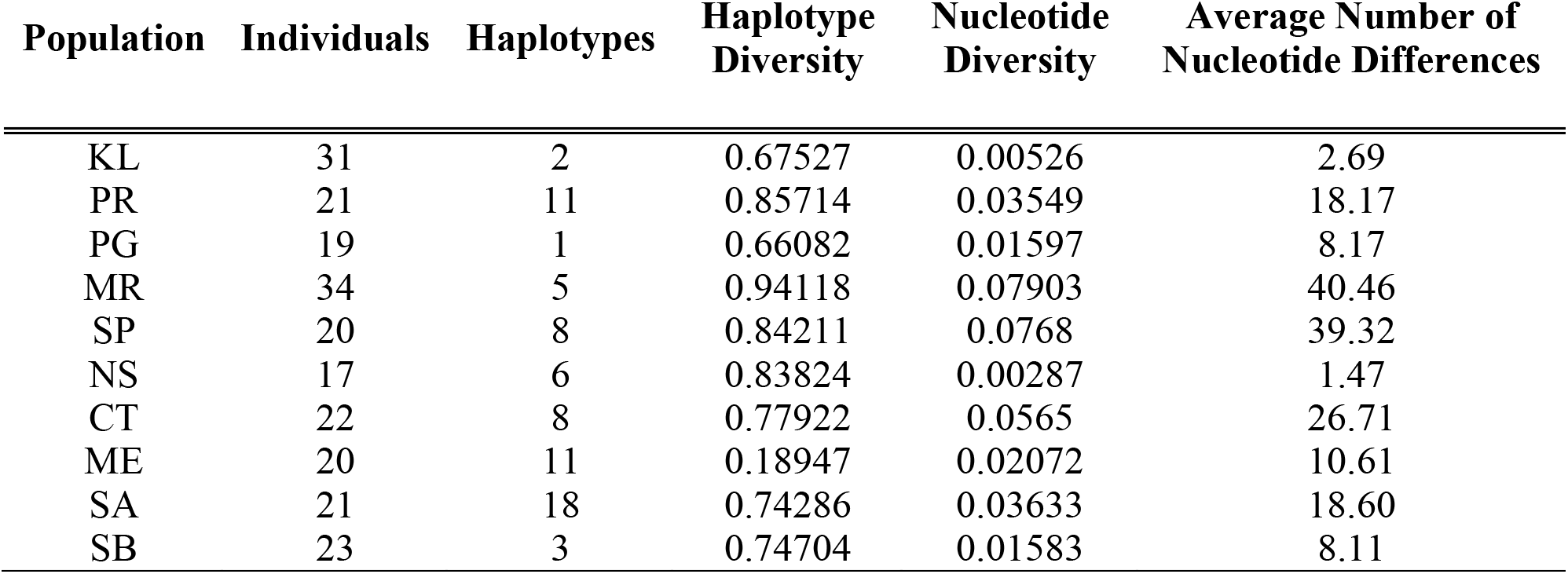
Population genetic summary statistics, as calculated by DNaSP.

| Population | Individuals | Haplotypes | Haplotype Diversity | Nucleotide Diversity | Average Number of Nucleotide Differences |
| --- | --- | --- | --- | --- | --- |
| KL | 31 | 2 | 0.67527 | 0.00526 | 2.69 |
| PR | 21 | 11 | 0.85714 | 0.03549 | 18.17 |
| PG | 19 | 1 | 0.66082 | 0.01597 | 8.17 |
| MR | 34 | 5 | 0.94118 | 0.07903 | 40.46 |
| SP | 20 | 8 | 0.84211 | 0.0768 | 39.32 |
| NS | 17 | 6 | 0.83824 | 0.00287 | 1.47 |
| CT | 22 | 8 | 0.77922 | 0.0565 | 26.71 |
| ME | 20 | 11 | 0.18947 | 0.02072 | 10.61 |
| SA | 21 | 18 | 0.74286 | 0.03633 | 18.60 |
| SB | 23 | 3 | 0.74704 | 0.01583 | 8.11 |

In all, 73 unique haplotypes were identified, which segregated into four major clades (Fig. 8). Each clade was represented in populations from across a large geographic area, and most populations contained individuals from multiple clades. Additionally, we observed several instances of shared haplotypes between geographically distant populations (highlighted in Fig. 8). As exceptions to this, the Key Largo, Punta Gorda, Manatee River, and Shediac Bay populations, each of which were characterized by differentiated thermal performance curves, were all largely represented by a single dominant haplotype each, which was generally not recovered from other populations. The dominant single haplotype is, however, shared between Punta Gorda and Manatee River. Estimates of the number of migrants from MigrateN varied greatly between populations, ranging between 2.6 to 107.7 per generation. There was a negative correlation between the number of migrants exchanged and the pairwise population difference in LD_50_ values (p = 0.045; Supp. Fig. 6). However, this correlation explained only a very small proportion of the variance (R^2^ = 0.07).

**Figure 8:**
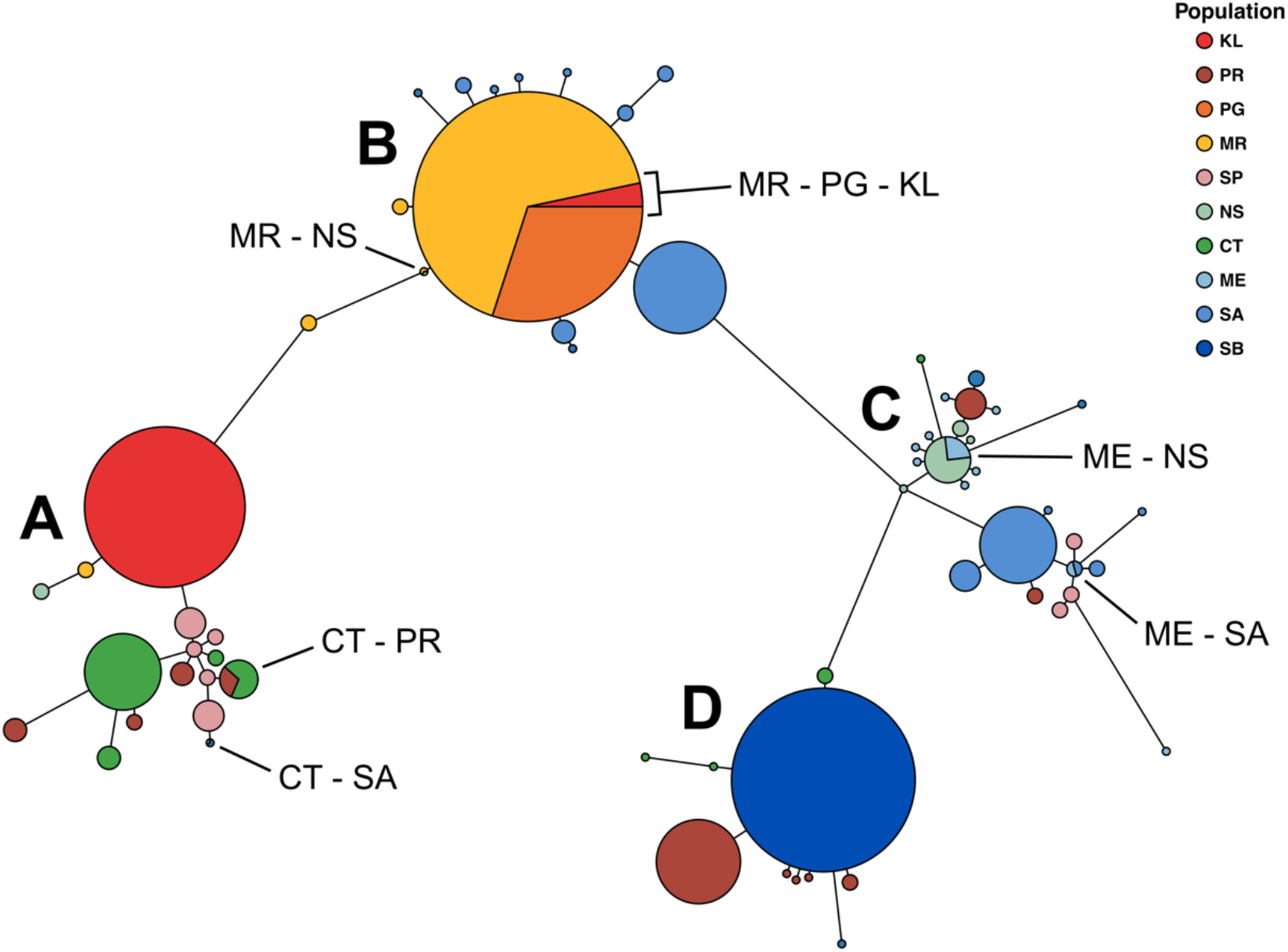
Mitochondrial COI haplotype network for *Acartia tonsa* sequences recovered from each of the ten populations. Individual circles represent unique haplotypes, with the size of each proportional to the haplotype’s frequency. Connecting bar lengths are proportional to the number of base pair differences between haplotypes. Circles are colored based on the sequence population of origin. Shared haplotypes are labelled with the sampling sites they were recovered from. Four distinct clades are observed and labelled A – D.

## Discussion

Thermal tolerance and the strength of phenotypic plasticity both vary strongly among populations of *Acartia tonsa* from across the North Atlantic and Gulf of Mexico. This variation conforms to the predictions of the CVH (Janzen, 1967; Stevens, 1989), but may also be explained by Stillman’s hypothesis regarding the trade-off between thermal tolerance and the strength of phenotypic plasticity (Stillman 2003). We also observed a surprising lack of differentiation of thermal performance across a large portion of the sampled range. Mitochondrial COI sequences suggest that this lack of differentiation is due to constraint by gene flow, rather than selection for a generalist performance curve. The patterns we observed in both thermal tolerance and the strength of phenotypic plasticity may result in regional differences in vulnerability to warming, with low latitude populations being the most vulnerable.

### Patterns in Thermal Adaptation

Key Largo (KL), the southernmost population, had the highest thermal tolerance of the ten populations examined, followed by three populations from the Gulf of Mexico. The northernmost population, Shediac Bay (SB), had the lowest thermal tolerance but the largest strength of phenotypic plasticity. Manatee River (MR), one of the Gulf of Mexico populations with higher thermal tolerance, had the smallest strength of phenotypic plasticity. The CVH (Janzen, 1967; Stevens, 1989) predicts higher thermal tolerance in warmer environments and increased phenotypic plasticity in more variable environments. The results of our study are consistent with these predictions. We observed significant correlations between I) thermal tolerance and mean monthly temperature and II) the strength of phenotypic plasticity and mean monthly temperature variance. The observed patterns, however, may also be explained by a trade-off between thermal tolerance and phenotypic plasticity (Stillman, 2003). We observed a significant negative relationship between thermal tolerance and the strength of phenotypic plasticity, but no significant correlation between mean monthly temperature and mean monthly temperature variance. Together, these observations might indicate that the trade-off proposed by Stillman (2003) has a strong influence on shaping patterns of adaptation in this taxon, as there is no covariance between environmental parameters that would drive the observed correlation between LD_50_ and ΔLD_50_. The few studies that have estimated the strength of selection on phenotypic plasticity generally indicate relatively weak selection (Arnold *et al*., 2019). It is not implausible, therefore, that observed spatial patterns in phenotypic plasticity might instead be driven by patterns of selection for increased thermal tolerance. However, support for this hypothesis appears to be limited and species specific (Gunderson & Stillman, 2015), warranting investigation in additional species.

The negative relationship between thermal tolerance and the strength of phenotypic plasticity might reflect a role of genetic accommodation or “plasticity first” evolutionary change (Scheiner *et al*., 2017; Kelly, 2019; Levis & Pfennig, 2016; Levis *et al*., 2017; Pfennig *et al*., 2010; Pigliucci *et al*., 2006; Price *et al*., 2003; West-Eberhard, 2005; 2003), wherein phenotypic modification by plasticity becomes canalized or fixed by genetic mechanisms, resulting in the loss of capacity for plastic change. The Manatee River - St. Petersburg (MR - SP) comparison best illustrates this in our study. These sites are in close proximity to each other (a feature discussed further below) but exhibit strong differences in both thermal tolerance and the strength of phenotypic plasticity. St. Petersburg had a lower thermal tolerance in the 18°C developmental temperature group, but larger strengths of phenotypic plasticity than Manatee River. Increasing developmental temperature reduces the difference in thermal tolerance between the two populations. If we assume that these two populations share an ancestral phenotype similar to the contemporary St. Petersburg population, the elevated thermal tolerance but reduced plasticity in Manatee River may reflect the fixation of changes originally induced by phenotypic plasticity in the Manatee River population. However, this is the only population pair where this is observed (the Punta Gorda (PG), Punta Rasa (PR), and Key Largo (KL) populations all had comparable or higher thermal tolerance values but larger strengths of phenotypic plasticity than the Manatee River population), suggesting that there may be several alternative evolutionary mechanisms at play across larger spatial scales.

### Spatial Scales of Adaptation - Gene flow vs. Selection

Local adaptation within the range of dispersal, so called microgeographic adaptation (Richardson *et al*., 2014), has garnered increased attention. “Microgeographic” may be a misleading term in marine systems though, as dispersal kernels can encompass hundreds of kilometers (Cowen & Sponaugle, 2009; Kinlan & Gaines, 2003; Kinlan *et al*., 2005). The lack of differentiation we observed over a large geographic area, ranging from the Gulf of Mexico to the Bay of Fundy, is in agreement with the expectation of high levels of gene flow in pelagic copepods and other planktonic taxa. The wide distribution of genetic clades, several instances of shared haplotypes between distant populations, and the negative correlation between number of migrants and the pairwise difference in LD_50_ suggests that gene flow can be strong enough to constrain the adaptive divergence of thermal performance curves in this taxon. Direct dispersal across these large distances within the short generation times of *A. tonsa* is unlikely. Instead, dispersal may follow a stepping-stone model (Kimura & Weiss, 1964), as is common in other coastal and estuarine taxa (Hellberg, 1995; Burridge *et al*., 2004; Williams *et al*., 2008; Ragionieri *et al*., 2010; Crandall *et al*., 2012).

In stark contrast to this pattern of long-distance connectivity, we also observed significant differentiation of TPCs over spatial scales of less than ten kilometers, suggestive of microgeographic adaptation. Copepods from the Manatee River (MR) and St. Petersburg (SP) sites were collected from either side of a strong salinity gradient, with the salinity at the St. Petersburg collection site approaching full oceanic levels (32 psu) and the Manatee River site strongly influenced by riverine input (5 psu). Adaptation to salinity has been shown to correspond with reproductive isolation in other populations of *Acartia tonsa* (Plough *et al*., 2018). In this case, dispersal and gene flow may be strongly limited by this salinity gradient, allowing for the local adaptation of the thermal performance curve in the Manatee River population, while the St. Petersburg population remains under the constraining influence of gene flow from other environments. This is supported by the population genetic data; the Manatee River individuals are almost entirely represented by a single haplotype in Clade B, while St. Petersburg individuals are represented by several different haplotypes in clades A, C, and D. No haplotypes are shared between these two sites, despite their distinct geographic proximity. It should also be noted that the major haplotype found at Manatee River represents the only haplotype found at Punta Gorda (PG), the other low salinity site included in this study. However, unlike in the Manatee River - St. Petersburg comparison, the Punta Gorda (PG) and the adjacent Punta Rasa (PR) populations share a similar TPC, despite being genetically distinct. The apparent local adaptation of both populations suggests that factors other than salinity may also contribute to isolation and the reduction of gene flow between Punta Rasa and other sites.

Interestingly, differentiation of TPCs occurs predominantly within clades rather than between clades, suggesting a rapid rate of thermal adaptation relative to the differentiation of the COI marker region. Each clade contains several distinct thermal phenotypes. Clade A for example contains populations representing three of the distinct groups of TPCs. This appears to be true for both temperature and salinity conditions, as clade B is recovered from warm and fresh sites like Manatee River (MR) and Punta Gorda (PG) as well as cold and high salinity sites like St. Andrew (SA). The generally large capacity for plasticity observed in *Acartia tonsa* may promote this wide distribution of clades across environmental conditions. Phenotypic plasticity may allow migrants to survive in a wider range of environments, thus increasing gene flow and constraining local adaptation (Crispo, 2008; Thibert-Plante & Hendry, 2011).

Of course, as is the case for all single-gene markers studies, our results must be interpreted with caution. Single-gene markers are vulnerable to incomplete lineage sorting (Nichols, 2001), and may obscure important patterns in genetic structuring. Genome-scale data would provide many more markers for population genomic assessment of structure and gene flow, but for the purposes of this study COI sequences provide robust evidence for where, across the examined range, gene flow may be potentially restricting adaptive divergence of performance curves.

### Vulnerability to Climate Change

Because copepods are key components in marine ecosystems and biogeochemical cycles determining their susceptibility to warming is essential for predicting the fate of the oceanic biota in light of climate change. An increasingly large body of literature has recognized the important role phenotypic plasticity may play in determining organismal responses to rapid climate change (Somero, 2010). Much of this literature has focused on acclimation or hardening, but our results show clearly that developmental phenotypic plasticity also deserves increased scrutiny as a factor affecting vulnerability (Burggren, 2018). In most of the populations examined, we observed an almost 1:1 relationship between the increase in thermal tolerance compared with the increase in developmental temperature. Strong phenotypic plasticity like this may reduce vulnerability to rapid climate change by providing a mechanism for correspondingly rapid phenotypic change. However, developmental phenotypic plasticity had a weaker effect on thermal limits than thermal tolerance (ΔLD_10_ < ΔLD_50_; Supp. Fig. 1). The apparent constraint of upper thermal limits suggests that accommodation of future warming by plasticity alone may increase vulnerability to extreme temperature events like heatwaves (Meehl, 2004; Perkins *et al*., 2012), as the difference between environmental temperature and thermal limits decreases.

In addition to strong phenotypic plasticity, the high levels of genetic diversity, a large potential for gene flow, and the apparently rapid rate of thermal adaptation (when gene flow is weak) would also suggest reduced vulnerability to warming. Selection on standing genetic variation may provide a rapid response to change (Pantel *et al*., 2015; Barrett & Schluter, 2008; Torda *et al*., 2017). Plasticity may also play a role in increasing or maintaining migration between populations, indirectly supporting evolutionary rescue (Crispo, 2008). Paired with the high levels of standing genetic diversity observed both within and between populations, increasing migration success may reduce vulnerability. This makes understanding oceanographic effects of climate change an important prerequisite for predicting biotic responses, as changes in ocean current patterns may strongly affect the potential for gene flow between populations. Reductions in gene flow may promote local adaptation, thus reducing vulnerability, while an increase in gene flow could strongly increase vulnerability to climate change if existing local adaptation is eroded by gene swamping (Lenormand, 2002).

Vulnerability will also be affected by the pre-existing spatial patterns in adaptation. Using the current and predicted temperatures at each site, we observed large variation in the potential change in survivorship at maximum temperatures. In the southern range, warming generally had little effect on survivorship at the maximum temperature in locally adapted populations, likely due to accommodation by developmental phenotypic plasticity. While the response was mixed, non-differentiated populations (those constrained by gene flow) may be strongly affected by warming; the St. Petersburg (SP) population saw the largest decrease in survivorship. Both non-differentiated and locally adapted populations from higher latitudes generally saw increased survivorship. Stronger predicted warming in this region may drive a larger increase in thermal tolerance due to developmental phenotypic plasticity, especially in the northernmost population Shediac Bay (SB), which had the largest strength of phenotypic plasticity. In general, local adaptation to increased temperature does not appear to increase vulnerability to warming, contrary to what has been previously suggested for warm-adapted tropical species (Somero, 2010; Tewksbury *et al*., 2008; Nguyen *et al*., 2011). However, this is likely highly regionally specific; the second largest decrease in survivorship was predicted for one of the locally adapted populations, Punta Rasa (PR), from the Gulf of Mexico.

Our analysis incorporates population differences in both thermal tolerance and the strength of phenotypic plasticity to provide a more robust estimate of vulnerability. Previous work also indicated that warm-adapted low latitude populations may be more vulnerable to warming, as they already experience temperatures near their thermal limits (Sasaki *et al*., 2019). This study refines that prediction with the inclusion of more populations from a wider range of thermal environments, and the integration of population genetic data. However, our analysis cannot account for other important factors that will also likely play a large role in determining vulnerability to climate change. Factors such as changing food quality and quantity (Gregg *et al*., 2003; Van der Waal *et al*., 2010; Paul *et al*., 2015; Hixon & Arts, 2016; Dutkiewicz *et al*., 2019), changes in predation pressure (Broitman *et al*., 2009; Rall *et al*., 2009; De Block *et al*., 2013; Allan *et al*., 2015), phenological mismatches between copepods, their prey, and their predators (Edwards & Richardson, 2004; Søreide *et al*., 2010; Brown *et al*., 2016), changes in behavior (Kearney *et al*., 2009; Marshall *et al*., 2013; Nagelkerken & Munday, 2015), and changing direction or magnitude of gene flow between populations all might strongly shape population vulnerability to climate change. Our analysis also assumes that populations are not undergoing range shifts or further evolutionary adaptation. Despite these limitations, our results provide a representative baseline estimate of vulnerability incorporating several different adaptive mechanisms.

The major conclusions of this study are possible only through the integration of physiological experiments and molecular ecology. On their own, the thermal performance curves cannot differentiate the potential explanations for the lack of divergence observed across large distances, selection for a generalist performance curve or gene flow over evolutionary timescales. The population genetic insights alone are not enough to infer vulnerability. We demonstrate that tight integration between different fields can provide a more comprehensive understanding of the factors determining vulnerability to climate change.

## Acknowledgements

We thank Dr. Ann Bucklin for helpful comments on the analysis of the molecular data, and Dr. Kendra Daly for assistance sampling in St. Petersburg. Research was supported by grants NSF-OCE 1559180 and CT Sea Grant R/LR-25, a Research Council grant from the University of Connecticut, and graduate research fellowships from the Department of Marine Sciences, University of Connecticut, USA.

## Supplementary Table

**Supp. Table 1:**
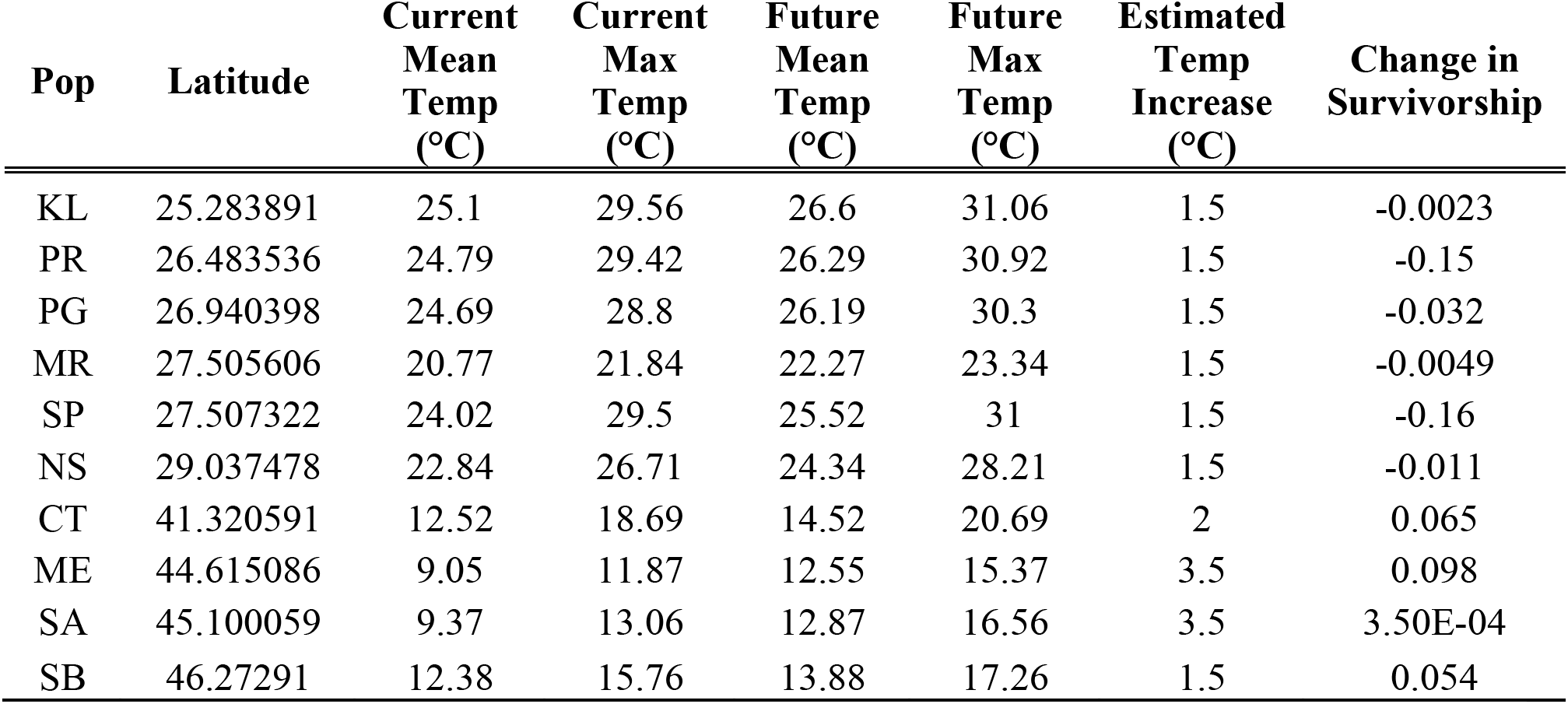
Temperature data used for predictions of vulnerability to warming. Current temperature data was acquired from the MODIS-aqua SST database. Warming was estimated from a high-resolution model of warming in the North Atlantic (Saba *et al*. 2016). Change in survivorship at maximum temperatures was estimated using TPCs based on a developmental temperature equal to the mean temperature.

## Supplementary Figures

**Supp. Fig. 1:**
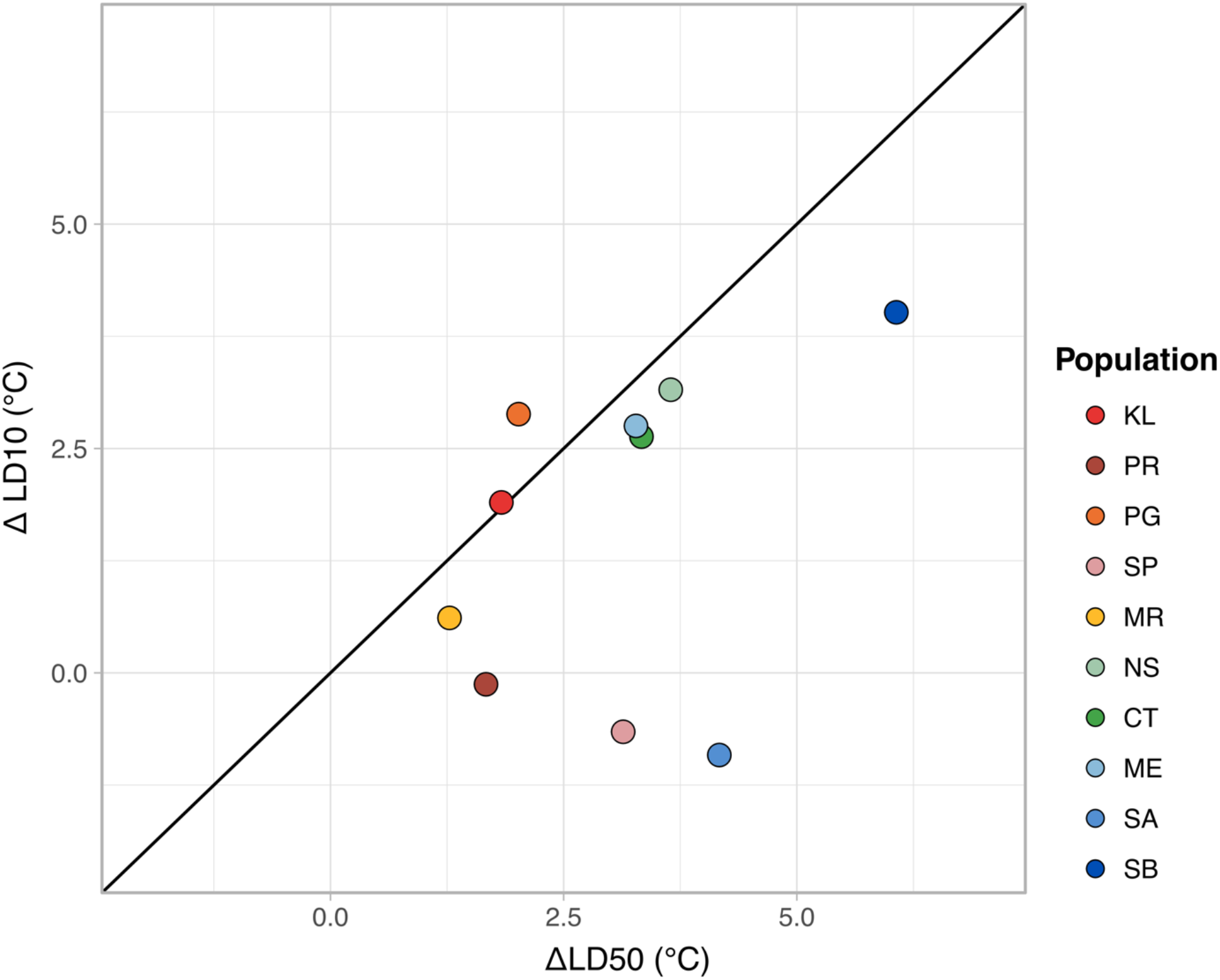
Change in thermal limits vs. change in thermal tolerance by developmental phenotypic plasticity. The solid line represents a 1:1 relationship. Points that fall below the line represent a larger increase in thermal tolerance than thermal limits.

**Supp. Fig. 2:**
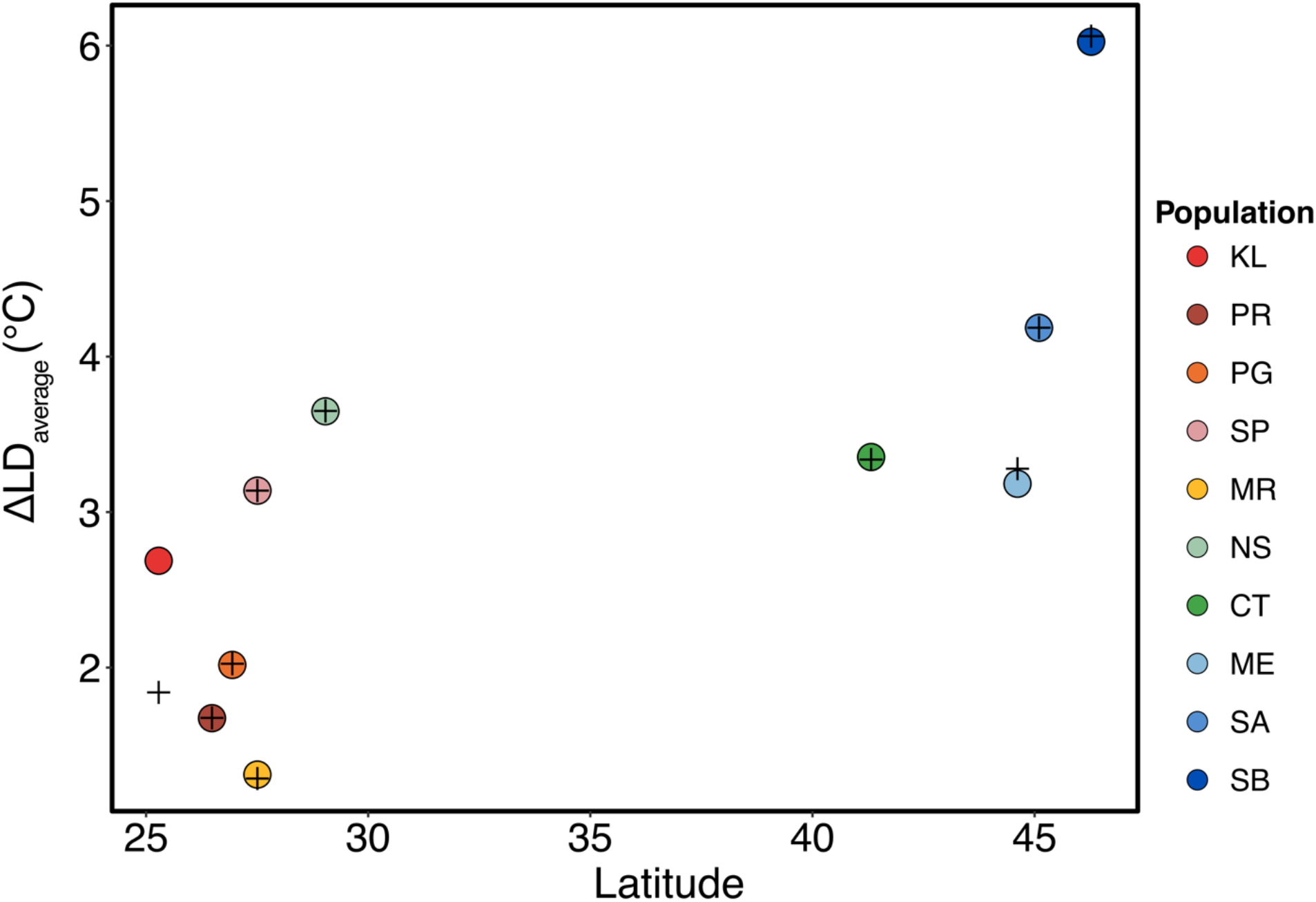
Average increase in survivorship at every dosage level. No major differences are observed between this metric and ΔLD_50_ values, which are represented by black crosses.

**Supp. Fig. 3:**
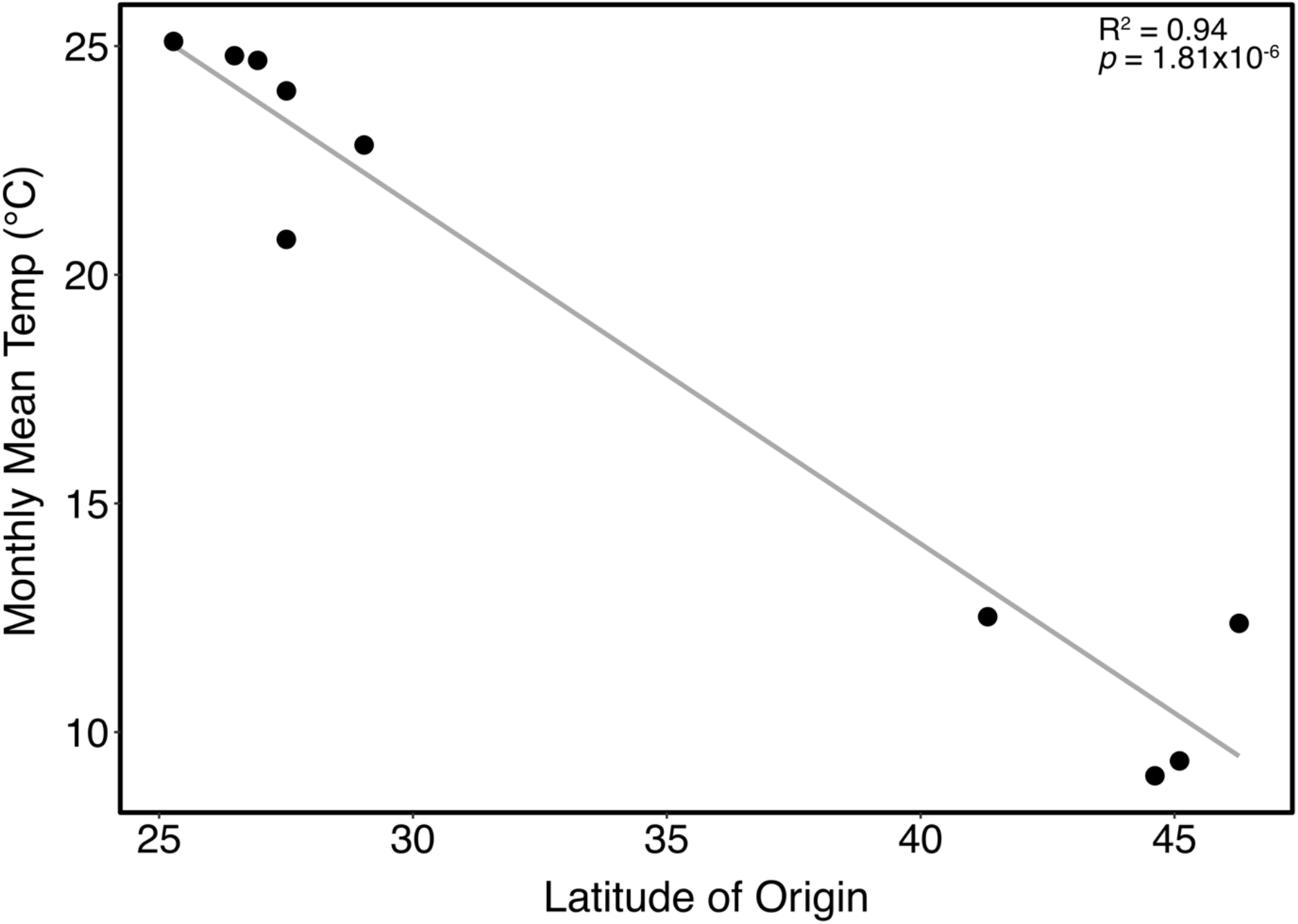
Correlation between monthly mean temperature and latitude. A significant correlation is observed.

**Supp. Fig. 4:**
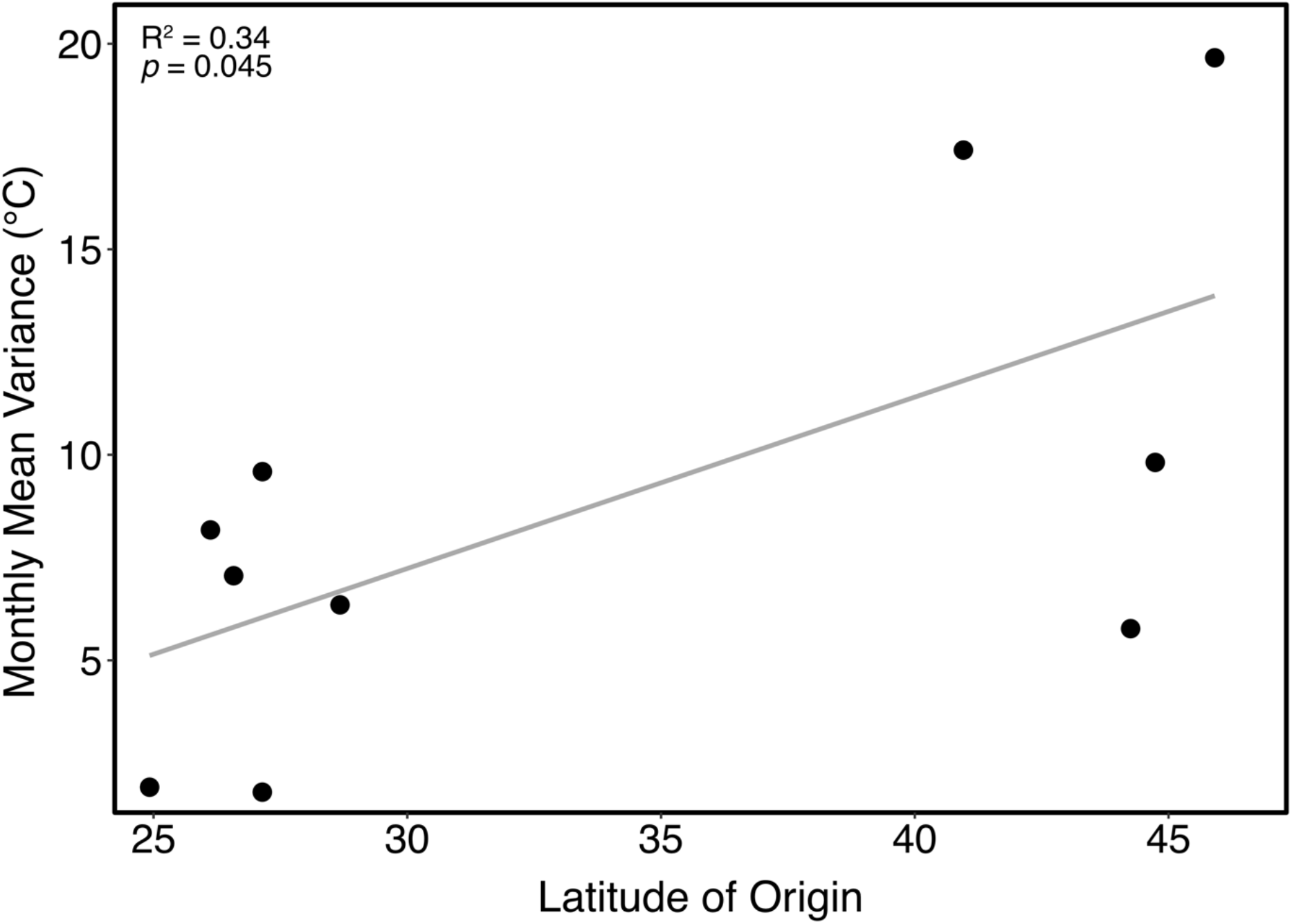
Correlation between monthly mean temperature variance and latitude. A significant correlation is observed.

**Supp. Fig. 5:**
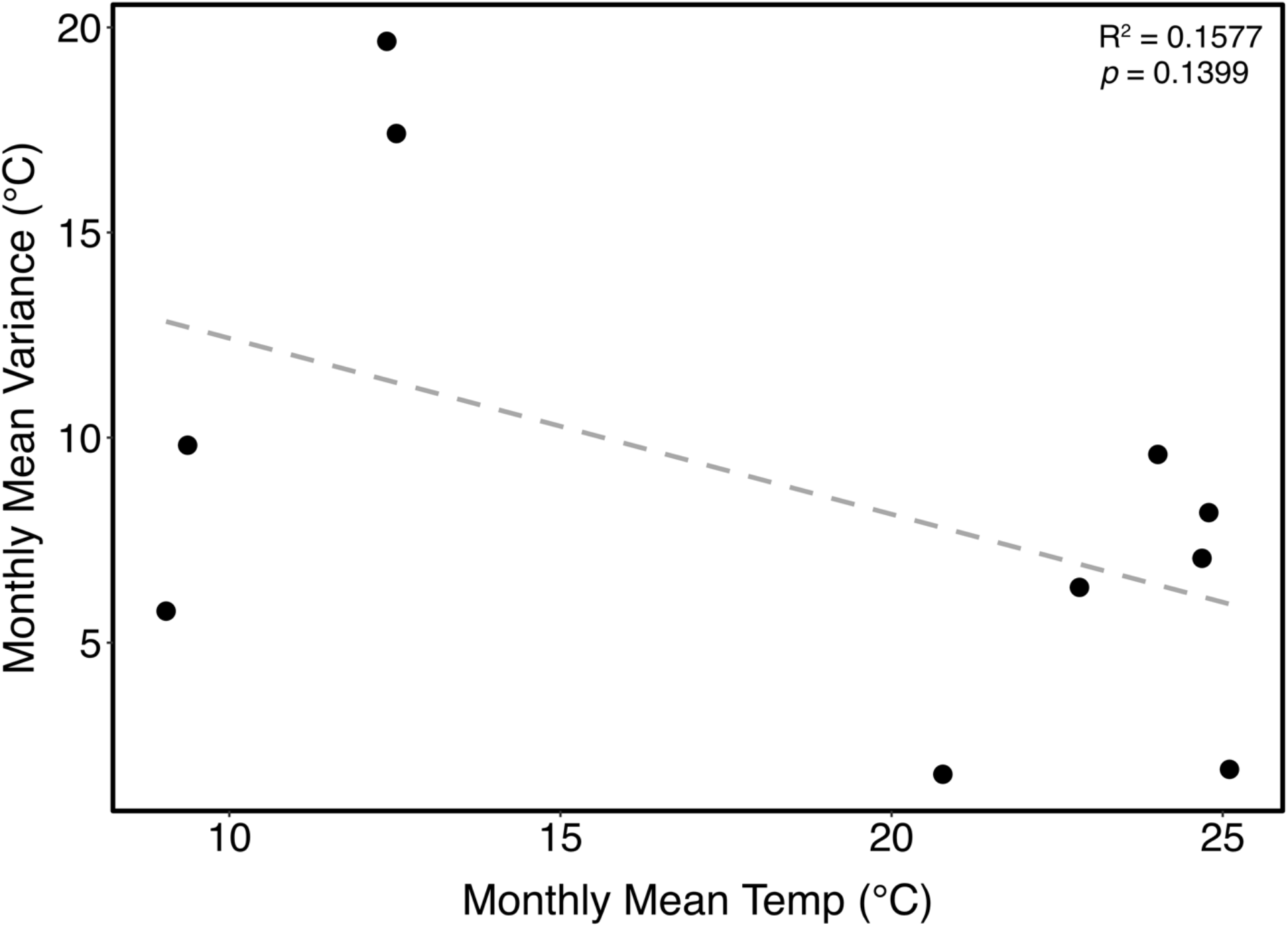
Correlation between the two explanatory environmental variables (mean monthly temperature and mean monthly temperature variance). No significant relationship between the variables is observed.

**Supp. Fig. 6.**
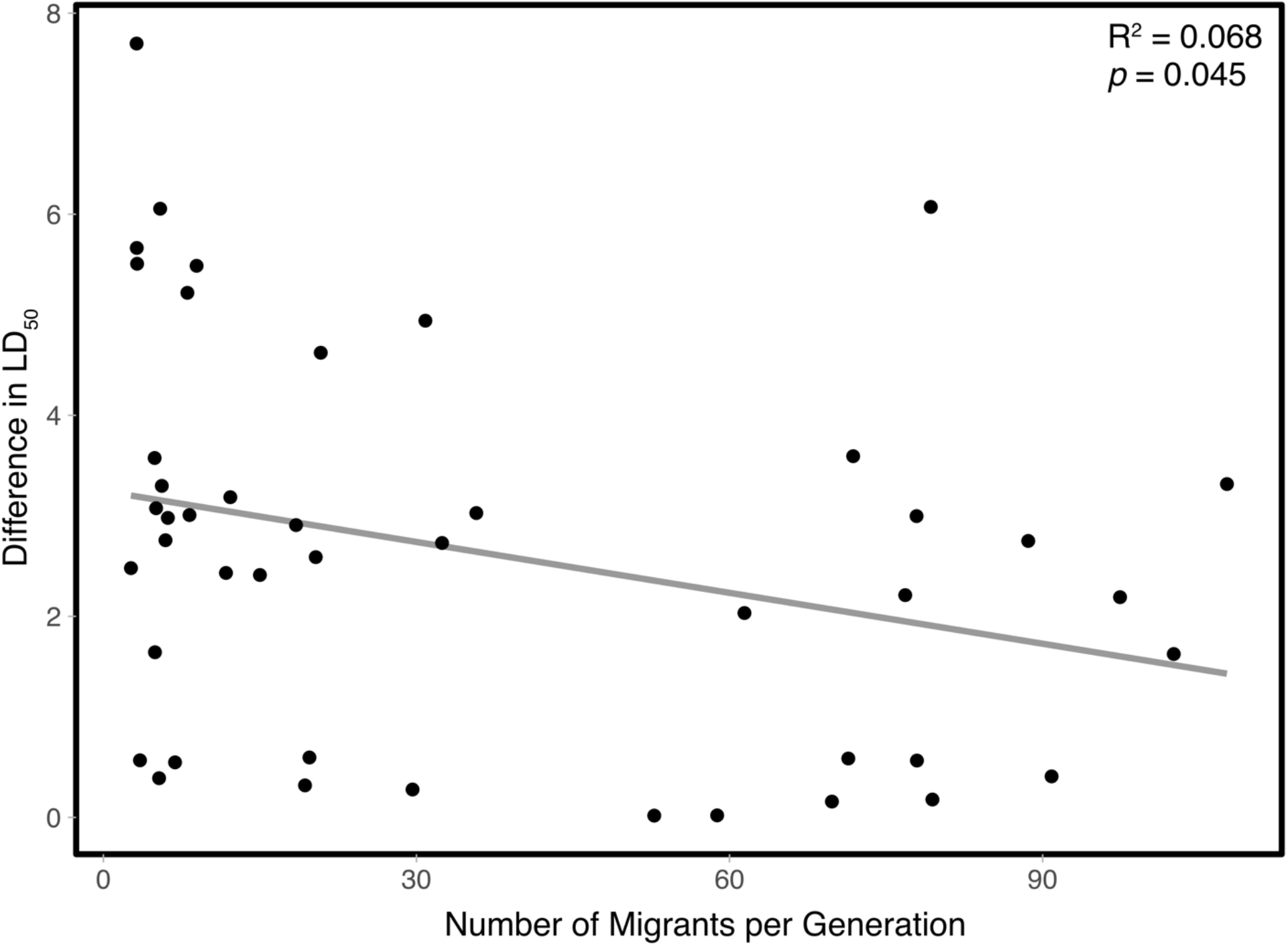
Correlation between the estimated number of migrants exchanged between two populations and the corresponding pairwise difference in thermal tolerance values between those populations.

